# *Acinetobacter baumannii* DacC influences cell shape, biofilm formation and physiological fitness by manifesting DD-carboxypeptidase, and β-lactamase dual-enzyme activities

**DOI:** 10.1101/2024.07.16.603720

**Authors:** Shilpa Pal, Diamond Jain, Sarmistha Biswal, Sumit Kumar Rastogi, Gaurav Kumar, Anindya S Ghosh

**Affiliations:** Department of Bioscience and Biotechnology, Indian Institute of Technology Kharagpur, West Bengal, India, PIN-721302

**Author notes:** Correspondence: Anindya S. Ghosh **Contact Information** Department of Biotechnology, Indian Institute of Technology Kharagpur, West Bengal PIN-721302, India.

**Keywords:** *Acinetobacter baumannii*, Penicillin-binding-protein (PBP), DD-Carboxypeptidase, β-lactamase, cell shape

## Abstract

With the growing threat of drug-resistant *Acinetobacter baumannii*, there is an urgent need to comprehensively understand the physiology of this nosocomial pathogen. As penicillin-binding proteins are attractive targets for antibacterial therapy, herein we have tried to explore the physiological roles of two putative DD-carboxypeptidases, viz., *dacC* and *dacD* in *A. baumannii*. Surprisingly, the deletion of *dacC* resulted in a reduced growth rate, loss of rod-shaped morphology, reduction in biofilm-forming ability, and enhanced susceptibility towards β-lactams, whereas, the deletion of *dacD* had no such effect. Interestingly, ectopic expression of *dacC* restored the lost phenotypes. The double deletion mutant in which both *dacC* and *dacD* were absent showed properties similar to the *dacC* single knockout. On the other hand, cell-shape reverting efficiency in septuple PBP deleted *E. coli* and *in vitro* enzyme kinetics assessments reveal that *dacD* is a stronger DD-CPase as compared to *dacC*. The expression of *dacC* was in the log phase whereas *dacD* expression takes place in the stationary phase. In summary, we conclude that *dacC* encodes a dual enzyme, possessing activities of DD-CPase and β-lactamase, which significantly affects the physiology of *A. baumannii* in various ways whereas *dacD* encodes a strong DD-CPase and plays a role in cell morphology, though it exerts negligible impact on other physiological aspects like intrinsic antibiotic resistance or biofilm formation.

**Graphical Abstract:** 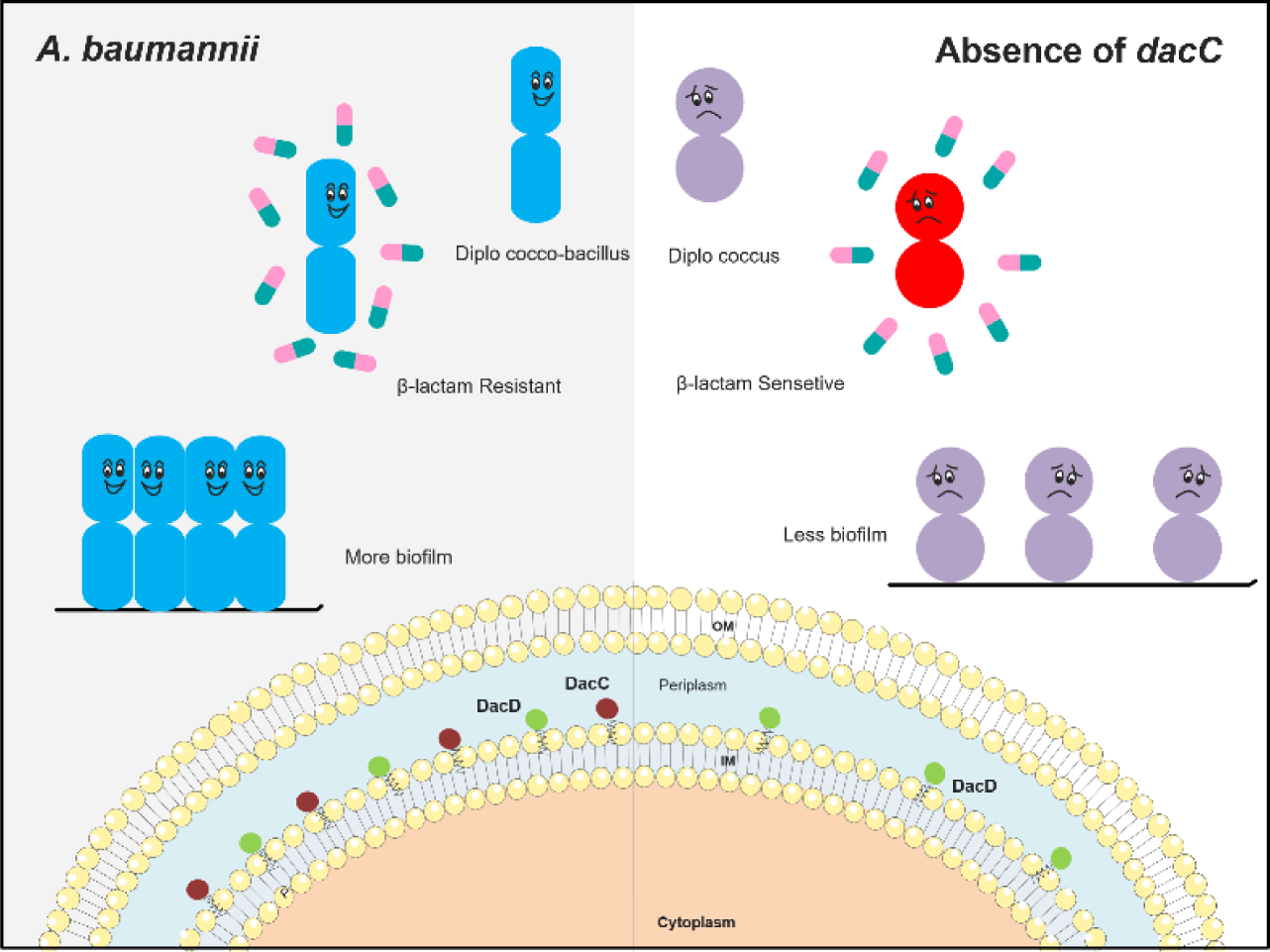

## Introduction

Nosocomial infections have become a great challenge for modern medicine. The leading pathogens of nosocomial infections include species of *Enterococcus faecalis*, *Staphylococcus aureus*, *Klebsiella pneumoniae*, *Acinetobacter baumannii*, *Pseudomonas aeruginosa*, and *Enterobacter spp*. (abv. ESKAPE). These ‘ESKAPE’ pathogens are so named to emphasize their ability to ‘escape’ the effects of antimicrobial treatment due to acquisition of resistance genes as well as formation of biofilms.

*Acinetobacter baumannii*, is an encapsulated Gram-negative aerobic coccobacillus found in soil and water (Van Looveren et al., 2004). In 1970s it was susceptible to common antibiotics, but in the last decade this opportunistic pathogen has emerged as one of the most common and deadliest ‘Superbugs’ and has become one of the prime causes of hospital acquired infections across the globe. They manifest serious diseases by frequently infecting immuno-compromised human hosts, like ventilator-acquired pneumonia, wound infections, urinary tract infections, septicaemia, meningitis, and necrotizing fasciitis resulting in significant morbidity and mortality. With the worldwide emergence of multidrug resistant strains, there is an urgent need to identify new drug targets (Karageorgopoulos and Falagas, 2008; Beceiro et al., 2014).

Elucidation of the mechanism of resistance by pathogens is the first step towards the development of novel approaches for the treatment and prevention of infections and for designing future chemotherapeutics. Peptidoglycan (PG) is still considered as an attractive target in chemotherapeutics because of its absence in the host cells, thereby making the selectively toxic drugs that target PG synthesis or modulation, specific for bacteria. The bacterial peptidoglycan, a three-dimensional, net-like mesh which lines the exterior of the cell membrane, is made up of glycan chains of alternating N-acetylglucosamine and N-acetylmuramic acid, cross-linked by short stem peptides attached to the N-acetylmuramic acid (Sauvage et al., 2008). The enzymes that are involved in peptidoglycan synthesis and remodelling are called Penicillin-Binding Proteins (PBPs), which are of two types, namely high molecular mass (HMM) and low molecular mass (LMM) PBPs. HMM PBPs catalyse the polymerization of the glycan strands (transglycosylation) and form the cross-links between glycan chains (transpeptidation) whereas LMM PBPs facilitate PG hydrolysis by catalysing the hydrolysis of the last D-alanine of stem pentapeptides (DD-carboxypeptidation) or by hydrolysing the peptide bond connecting two glycan strands (endopeptidation). The latter two classes of PBPs are called peptidoglycan remodelling enzymes, of which DD-carboxypeptidases have utmost importance (Ghosh et al., 2008). As their name suggests, PBPs bind to and are the targets of β-lactam antibiotics. Because of the structural resemblance between their natural substrate, the D-Ala-D-Ala end of the stem pentapeptide precursors and penicillin, the late-stage peptidoglycan synthesizing enzymes are sensitive to penicillin with which they form a long-lived acyl-enzyme that impairs their ability to crosslink peptidoglycan. PG remodelling enzymes, especially DD-carboxypeptidases, do have multiple functions as revealed by the various works on *E. coli* that include maintaining cellular morphology (Nelson and Young, 2000; 2001a; Ghosh and Young, 2003) providing intrinsic beta-lactam resistance (Sarkar et al., 2010; Sarkar et al., 2011), capable of showing beta-lactamase like activity upon single amino acid substitution (Dutta et al., 2015a; Dutta et al., 2015b; Kar et al., 2018) and influence biofilm formation (Gallant et al., 2005).

Though, there are multiple genes coding for different DD-CPases, they exhibit functional variation among different bacterial species (Ghosh and Young, 2003; Ghosh et al., 2008). In *A. baumannii*, three putative DD-CPases are annotated, *viz.*, PBP5/6 (gene *dacC*), PBP6b (*dacD*) and PBP7*(pbpG*) (Cayô et al., 2011). The deletion of *dacC* resulted in the loss of rod-shape morphology of *A. baumannii*, while PBP7 is proposed to possess DD-carboxypeptidase and DD-endopeptidase activities (Geisinger et al., 2020; Islam et al., 2022). Here, with the help of molecular genetics, biochemical and physiological studies, we attempted to elucidate the role of the DD-carboxypeptidase PBP5/6 *(dacC)* and the putative DD-CPase PBP6b *dacD* on the cell shape, antibiotic susceptibility, and biofilm formation, which to our knowledge might help designing future therapeutics.

## Material and Methods

### Bacterial strains, plasmids, and culture conditions

The bacterial strains and plasmids used here are listed in **Table 1**. *A. baumannii* ATCC 19606 from ATCC (Manassas, VA, USA). Bacterial cultures were grown at 37 °C in LB broth or LB agar (Himedia Mumbai, India) with appropriate antibiotics [Ampicillin (Amp), 100 µg mL^-1^; Chloramphenicol (Cam), 30 µg mL^-1^; Kanamycin (Kan), 50 µg mL^-1^; Tetracycline (Tet), 20 µg mL^-1^]. Antibiotic susceptibility testing was performed in cation-adjusted Mueller-Hinton broth (CAMHB) according to the CLSI guidelines (CLSI, 2012). Unless otherwise specified, all the molecular biology enzymes were purchased from New England Biolabs (Ipswich, MA, USA) and all the reagents from Sigma Chemical Co. (St. Louis, MO, USA). The molecular biology kits were purchased from Qiagen Inc. (Venlo, The Netherlands) and the manufacturer’s protocol was followed. Unless otherwise stated, routine DNA manipulations were performed by using standard procedures.

**Table 1.**
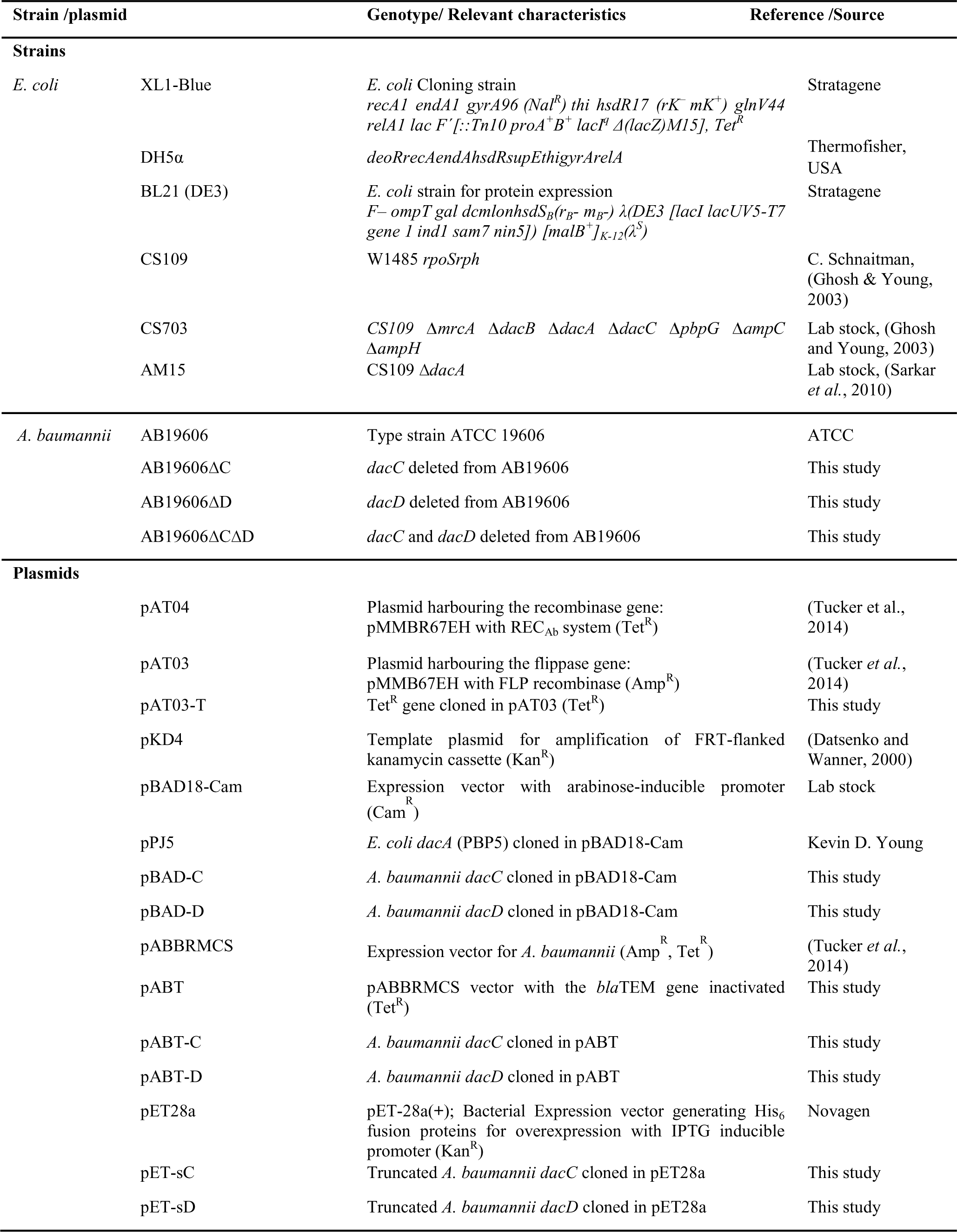
Strains and Plasmids.

### Construction of *A. baumannii* deletion mutants

The single and double deletion mutants of *dacC* and *dacD* were created in *A. baumannii* AB19606 (Cayô et al., 2011) by following the method described earlier (Tucker et al., 2014). In brief, *A. baumannii* strains carrying the plasmid pAT04 (harbouring the recombinase gene *rec_Ab_*) were grown at 37 °C for 45 mins at which the recombinase gene was induced using 2 mM IPTG and the cells were grown upto OD_600_ ∼0.4. The cells were harvested and electro-competent cells were prepared. The kanamycin cassette (Kan^R^) was amplified from the plasmid pKD4 along with the FRT (flippase recognition target) sites on either side, such that the amplicon was flanked on both the sides by the portions of sequences adjacent to the gene to be deleted. Primers containing 125-bp homology flanking the gene of interest were used to amplify the PCR products (see **Table S1)**. The cells were transformed with the linear PCR product. After growing the cells for 4 h in LB containing 2 mM IPTG, the cells were plated on LB agar containing 25 µg mL^-1^ of kanamycin. The transformants were screened through PCR to confirm deletion (**Fig. S1a**). For the construction of the double deletion mutant AB19606ΔCΔD, the kanamycin cassette was cured from the strain AB19606ΔD. Briefly, electrocompetent cells of AB19606ΔD were transformed with the plasmid pAT03-T (plasmid harboring the flippase gene). The flippase gene was expressed by inducing the transformants with 2 mM IPTG in LB broth. The transformants that were sensitive to both kanamycin and tetracycline were selected. The single deletion mutant (devoid of any antibiotic marker) was transformed with pAT04 and the previously described procedure was repeated.

### Cloning of membrane bound forms of dacC and dacD of *A. baumannii* for *in vivo* expression

The membrane bound forms of the genes *dacC* (1149 bp) and *dacD* (1320 bp) were amplified using AB19606 genomic DNA as template by using sets of primers [**Table S1]** and cloned at *Nhe*I and *Kpn*I sites of the arabinose-inducible vector pBAD18-Cam (for expression in *E. coli*) and at the *Sac*I and *Kpn*I sites of the IPTG-inducible shuttle vector pABT (for expression in *A. baumannii*). The vector pABBRMCS (Tucker et al., 2014) has both ampicillin and tetracycline resistance cassettes. Therefore, to use the vector for *in vivo* expression studies for β-lactam susceptibility assay, the ampicillin resistance marker (*bla*TEM gene) was inactivated by mutating the Serine 68 to Alanine. The new vector had only one functional resistance marker and was named pABT. The sizes of the inserts corresponded to the size of the original genes (*dacC*-1149 bp, *dacD*-1320 bp) as confirmed by commercial sequencing service (Xceleris, Ahmedabad, India), and subsequently, the plasmids were introduced into the respective deletion mutants for *in vivo* complementation studies.

The vector pABT is IPTG inducible. Therefore, to finalize the concentration of IPTG to be used for *in vivo* expression studies with the complemented strains, we used the microbroth dilution method (used for antibiotic susceptibility studies). Briefly, the strains were inoculated in 96-well plates containing serial two-fold dilutions of IPTG in the liquid broth (experiments were performed with both LB and CAMH broth) and incubated overnight at 37 °C. After which the OD_600_ was measured to assess the growth of the cultures.

### Determination of growth curves

Growth curves of the *A. baumannii* strains were generated by inoculating the strains into 100 mL LB broth using 0.05% of inoculum from an overnight culture and incubating at 37 °C (without shaking) for 21 h. The optical density of cultures was measured at *A*_600_ every 30 min and the curves were plotted on a semilogarithmic base 2 scale (*y* axis).

### Semi-quantitative analysis of biofilm formation

Biofilm formation of the *A. baumannii* strains was assessed using the microtiter plate biofilm assay (Kumar et al., 2015). The wells of a sterile, 24-well polystyrene microtiter plate were filled with 1 mL LB and inoculated with ∼10^6^ cells. Following an incubation period of 24 h at 37 °C under static condition, the wells were washed with water to remove the planktonic cells and stained with 0.1% crystal violet (CV), and the excess CV was removed by washing with water. The bound dye was solubilized in 1 mL of 33% acetic acid, and the optical density of the acetic acid-dye solution was measured at *A*_620nm_ using an ultraviolet/visible spectrophotometer (Multiskan Spectrum-1500 Spectrophotometer).

### Microscopic analysis of the biofilms

Biofilm development was assessed qualitatively by doing the microscopic analysis using scanning electron microscopy (SEM). The biofilms were grown on cover-slips, fixed with 2% glutaraldehyde, dried and mounted on aluminium stubs. The samples were shadowed with gold and analyzed through a SEM (ZEISS EVO 60 Scanning Electron Microscope).

### Antibiotic susceptibility studies

Minimum Inhibitory Concentration (MIC) signifies the concentration of antibiotic at which the bacterial growth is visibly inhibited and it is a measure of the effectiveness of an antimicrobial agent. The effects of *dacC* and *dacD* on β-lactam resistance were assessed by determining the MIC of β-lactam antibiotics using the broth micro-dilution method following CLSI guidelines. Briefly, the assays were performed in microtitre plates with an assay volume of 300 μL of CAMHB and an inoculum size of ∼10^5^ cells per well. The strains harbouring recombinant plasmids were inoculated in the wells containing media supplemented with the respective inducers (arabinose/ IPTG) to induce the expression of the AB *dacC* and *dacD* clones. After incubating the plates at 37 °C for 16-18 h, the culture was measured at *A*_600nm_. All the experiments were repeated three times with three replicates in each set.

### Study of cell shape

The cell shape of *A. baumannii*, the deletion mutants and the complemented strains were analysed using bright field microscopy as described earlier (Nelson and Young, 2000). Cells were grown overnight in LB at 37 °C and following which was diluted to 0.01% and grown upto OD_600nm_∼0.4. Cells were harvested, washed with PBS and droplet of suspension was placed onto poly-lysine coated glass slides. Cells harbouring recombinant plasmids were induced at OD_600nm_∼0.2 with different concentrations of arabinose. Live cells were visualized using OLYMPUS IX71 microscope at 1000X magnification. The images were digitally captured using the cellSens software (Olympus Inc, Japan).

To assess the DD-CPase activities of dacC and dacD, the genes were expressed in *E. coli* CS703 cells (Ghosh and Young, 2003). CS703 cells containing pBAD18-Cam and pPJ5 (expressing *E. coli* PBP5 encoded by *dacA*) were used as negative and positive controls, respectively. Expression of PBP5 from pPJ5 was induced by using 0.0005% of arabinose, as higher inducer concentrations result in cell lysis (Nelson and Young, 2000). The cells were processed and visualized as mentioned above.

### RNA extraction and qPCR analysis

Overnight grown *A. baumannii* cells were sub-cultured in 100 mL of LB medium and grown at 37 °C. To collect cells from five different growth phases, namely, early log, mid log, late log, early stationary and late stationary, the cells were harvested at OD_600nm_ ∼0.2, 0.4, 0.6, 0.8 and 1, respectively. The total RNA was extracted using the RNeasy mini kit (Qiagen), and digested with DNase I (Thermo Scientific, USA) to remove the contaminating DNA. The corresponding cDNA was synthesized with Revert Aid H Minus First Strand cDNA synthesis kit (Thermo Scientific, USA) according to the manufacturer’s protocol. Real-time PCR reactions were performed in a volume of 25 μL using Maxima SYBR Green mix (Thermo Scientific, USA) in a 96-well plate. For relative quantification of gene expression, the comparative C_T_ method was used where the fold change was determined as 2^-ΔΔC^_T_. The 16SrRNA (housekeeping gene) was used as the endogenous control and the transcript expression level of the gene of interest was normalized to 16SrRNA levels. Fold changes are means ± SD and are derived from three independent RNA preparations. The qPCR primers used in this study were designed by Primer Express 3.0 (**Table S1**).

### Cloning, expression, and purification of the PBPs

Since, PBPs are membrane tethered proteins localised in the periplasmic space, therefore for the soluble expression of protein in the cytoplasm, the primers were designed in such a way that the amplicons were devoid of the signal peptide sequences and the C-terminal amphipathic anchors (**Table S1**). PBP5/6 has 382 amino acids (aa) and PBP6b has 439 aa. To solubilize DacC, 19 aa from the N-terminal and 15 aa from the C-terminal region were removed, resulting in a truncated protein of 348 aa. To solubilise DacD, 21 aa from the N-terminal and 30 aa from the C-terminal region were removed, and the resultant protein had 388 aa. The truncated PBP genes were amplified and cloned separately in the vector pET28a at the *Nhe*I and *Sac*I sites to create pET-C and pET-D **(Fig. S2d)** and confirmed by sequencing through commercial services (Xceleris, Ahmedabad, India).

For expression studies, the plasmids were transformed into *E. coli* BL21 cells and the conditions for protein expression were optimised. Bulk expression of sDacC was done by incubating the culture at 28 °C for 6 h post induction with 0.05 mM IPTG whereas sDacD was overexpressed at 16 °C for 16 h after inducing with 0.5 mM IPTG. The expressed proteins with N-terminal His-Tag were purified through Ni-NTA affinity chromatography using Akta Prime Protein Purification System (GE, USA). The proteins were eluted from the Ni-NTA column in a buffer (10 mM Tris-HCl, 300 mM NaCl buffer, pH 8.0) in the presence of imidazole at 4 °C. The proteins were dialysed for 16 h at 4 °C with three changes in the same buffer without imidazole. The dialysed proteins were subjected to 15% SDS-PAGE to determine their purity, and after estimating the concentrations, the PBPs were used for different assays. The molecular mass of purified proteins was determined via Matrix-Assisted Laser Desorption Ionization-Time of Flight (MALDI-Tof) spectra using VOYAGER-DE Pro, Applied Biosystem, USA. The mass spectra were acquired in the mass range of m/z 10000 to 40000 Da.

### in vitro assessment of DD-CPase activity

The DD-CPase activities of the proteins were determined by assessing the kinetics of the interaction of the proteins with peptide substrates. The peptidoglycan-mimetic pentapeptide, L-Ala-γ-D-Glu-L-Lys-D-Ala-D-Ala was used as substrate with concentrations varying from 2 mM to 10 mM in 50 mM Tris-HCl, pH 8.5, in the presence of the purified enzyme as illustrated previously (Chowdhury et al., 2010). D-Alanine produced in the reaction was measured and compared with standard D-alanine using a spectrophotometer (Multiskan Spectrum-1500 Spectrophotometer) at 460 nm. The data was analysed using the Enzyme Kinetic Module of SIGMAPLOT v13 (Systat Software) to determine the *k*_cat_ (turnover number) and the *K*_m_ (substrate concentration at half the maximum velocity).

### Kinetic analysis of the interaction of PBPs with Bocillin

The penicillin binding efficiency of the PBPs was assessed by their interaction with the commercially available fluorescent penicillin, BOCILLIN-FL (henceforth termed as Bocillin) (Zhao et al., 1999). The interaction was characterized by determining the rate of acyl–enzyme complex formation by calculating the second-order rate constant *k*_2_/*K* at sub-saturating concentrations of the substrate. The second-order rate constant *k*_2_/*K* is determined by calculating the pseudo-first-order rate constant *k*_a_ using the equation:

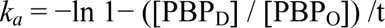

where ‘PBP_D_’ represents the density (experimentally derived) of the fluorophore associated with PBP bound Bocillin at time t for a particular Bocillin concentration D, and ‘PBPo’ shows the density at which the enzyme was saturated with Bocillin (Chowdhury et al., 2010, Chowdhury et al, 2012). The acylation rates of the PBPs were determined by incubating the enzymes with different concentrations (25, 50, 75 and 100 mM) of Bocillin in 10 mM phosphate buffer, pH 7.4 and incubating at 35 °C for various time intervals. The reactions were stopped by adding the sample buffer containing SDS and β-mercaptoethanol, and subsequently boiling the samples for 5 min. Bocillin-labelled protein samples were run through 15% SDS-PAGE and visualized under a Typhoon FLA 7000 (GE Healthcare, USA) at an excitation wavelength of 488 nm and an emission wavelength of 526 nm. The band intensities of the Bocillin bound proteins were measured by densitometric scanning (UVP Gel documentation system). Different *k_a_* values obtained were plotted against the corresponding Bocillin concentrations to determine *k_2_/K*.

The deacylation of the Bocillin-PBP complex results in the release of free PBP and the cleaved Bocillin. The deacylation reaction is described by the first-order rate constant *k*_3_, which is determined from the following equation:

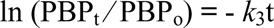

where, PBP_t_ is the residual acyl-enzyme concentration at time t, and PBP_o_ is the initial concentration of acyl-enzyme complex. The rate constant of the deacylation reaction was established by incubating the proteins with Bocillin (50 mM) for 15 min at 37 °C in 10 mM phosphate buffer, pH 7.4. An excess of Penicillin G was added and the amount of Bocillin remaining bound to the protein was determined by removing the aliquots at various time intervals. The intensity of labelled PBPs was measured by densitometric scanning as mentioned above. The deacylation rate constant (*k*_3_) was determined by measuring the fluorescence of the remaining PBP-bound Bocillin (the acyl–enzyme complex) that was reduced upon time, as described previously (Chowdhury et al., 2010, Chowdhury et al., 2012).

### Binding affinity analysis of the PBPs with β-lactams and inhibitors

The binding efficiency of the purified PBPs for different β-lactam antibiotics and inhibitors was determined by competition assays with Bocillin. The purified PBPs were incubated with increasing concentrations of each antibiotic/inhibitor for 20 min at 35 °C before Bocillin was added at a final concentration of 20 μM for an additional 20 min of incubation. The sample in which the PBPs were incubated with Bocillin and did not contain antibiotic was considered as 100% binding (Asli et al., 2016, Verma et al, 2023). The samples were run through SDS-PAGE and visualized as mentioned earlier. The fluorescence intensities of the Bocillin labelled PBPs were measured. The antibiotic concentration required to reduce the binding of Bocillin to the PBPs by 50% is referred as IC50. The values were estimated by plotting the PBP band intensities versus the antibiotic concentrations.

### Determination of the β-lactamase activity

The β-lactamase activities of the purified PBPs were determined by following a previously described method (Kumar et al., 2017). Briefly, the hydrolysis of different β-lactam antibiotics was monitored spectrophotometrically using a BioSpectrometer kinetic (Eppendorf Inc., Germany). The reactions were conducted by incubating the PBPs with the β-lactam substrates at 30 °C in 10 mM phosphate buffer (pH 7.4) and the changes in the absorbance was measured and plotted. The kinetic constants like enzyme affinity (*K*_m_), turnover number (*k*_cat_), and enzyme efficiency (*k*_cat_/*K*_m_) were obtained by determining the initial rate of the reaction at different substrate concentrations. The concentration-dependence of the initial rate was fitted and analysed using online curve-fitting tool (https://www.mycurvefit.com, My Assays Ltd., Brighton, UK). The concentrations of the substrates used, specific wavelengths for different antibiotics, and their absorption coefficients are tabulated in **Table S2.**

### Statistical Analysis

All the experiments were performed in triplicate, unless otherwise mentioned, and the results are expressed as mean ± standard deviation. The statistical significance (*P* value) of the data obtained was determined by performing the student’s t-test. A p-value is less than 0.05, is flagged with one asterisk (*), a p-value is less than 0.01 is flagged with two asterisks (**) and a p-value less than 0.001 is flagged with three asterisks (***). The experiments to determine MICs were performed in triplicate and repeated thrice and the most reproducible results were reported. The fold difference was calculated to represent the change in different parameters. It is defined as the ratio of values between the strains, provided the ratio ≥ 1 or, its reciprocal if the ratio is < 1. The IC_50_ for each antibiotic of both the PBPs were calculated from at least three independent PBP binding assays.

## Results

### Both DacC and DacD of A. baumannii reverse the morphological oddities of a septuple PBP deletion mutant to E. coli where the in vivo performance of AbDacD supersedes AbDacC

DD-CPases remove the terminal D-Ala from pentapeptides that control the levels of PG substrates available for the PG synthetic enzymes and preventing unwanted cross-link formation within the PG layer (Peters et al., 2016b). These enzymes are also involved in maintaining the cell shape of the host bacteria. *E. coli* CS703 is a seven (septuple) PBP deleted strain, rendering the cell deformed (Ghosh and Young, 2003; Dutta et al., 2015a). Ectopic expression of DD-CPases in *E. coli* CS703 can complement the shape back to normal (Nelson and Young, 2001b; Sarkar et al., 2011; Dutta et al., 2015a). Therefore, to understand the role of DacC and DacD in maintaining the cell shape, the corresponding genes were expressed ectopically in CS703 cells and checked for the aforementioned reversal of cell shape deformity.

The expression of both *dacC* and *dacD,* individually, resulted in the reversal of aberrant morphology of CS703 depending upon the concentration of inducer used for the expression. We observed that the overexpression of *dacD* could reverse the deformities of the CS703 upon induction with 0.0005% arabinose and the attempt to induce the expression of *dacD* with higher concentration of inducer (0.2% arabinose) resulted in a spherical cell shape that eventually led to cell lysis **(Fig. 1).** Overexpression of *dacC* could reverse the aberrations of CS703 at a higher inducer concentration (0.2% arabinose) (**Fig. 1**). These results indicate that both the PBPs can restore the normal cell shape of the deformed *E. coli* host, indicating the *in vivo* DD-CPase activity. Also, it has been reported that overexpression of strong DD-CPases result in the cell lysis (Nelson and Young, 2001a). As *AbdacD* required much lower concentration of arabinose than that of *AbdacC* to complement the cell shape oddities of *E. coli* CS703 *in vivo*, we speculated that AbDacD has higher *in vivo* DD-CPase activity as compared to ABDacC. Based on these observations, we attempted to study the effect of deletions of these two PBPs, either individually or in combination, on the cell shape of *A. baumannii*.

**Figure 1.**
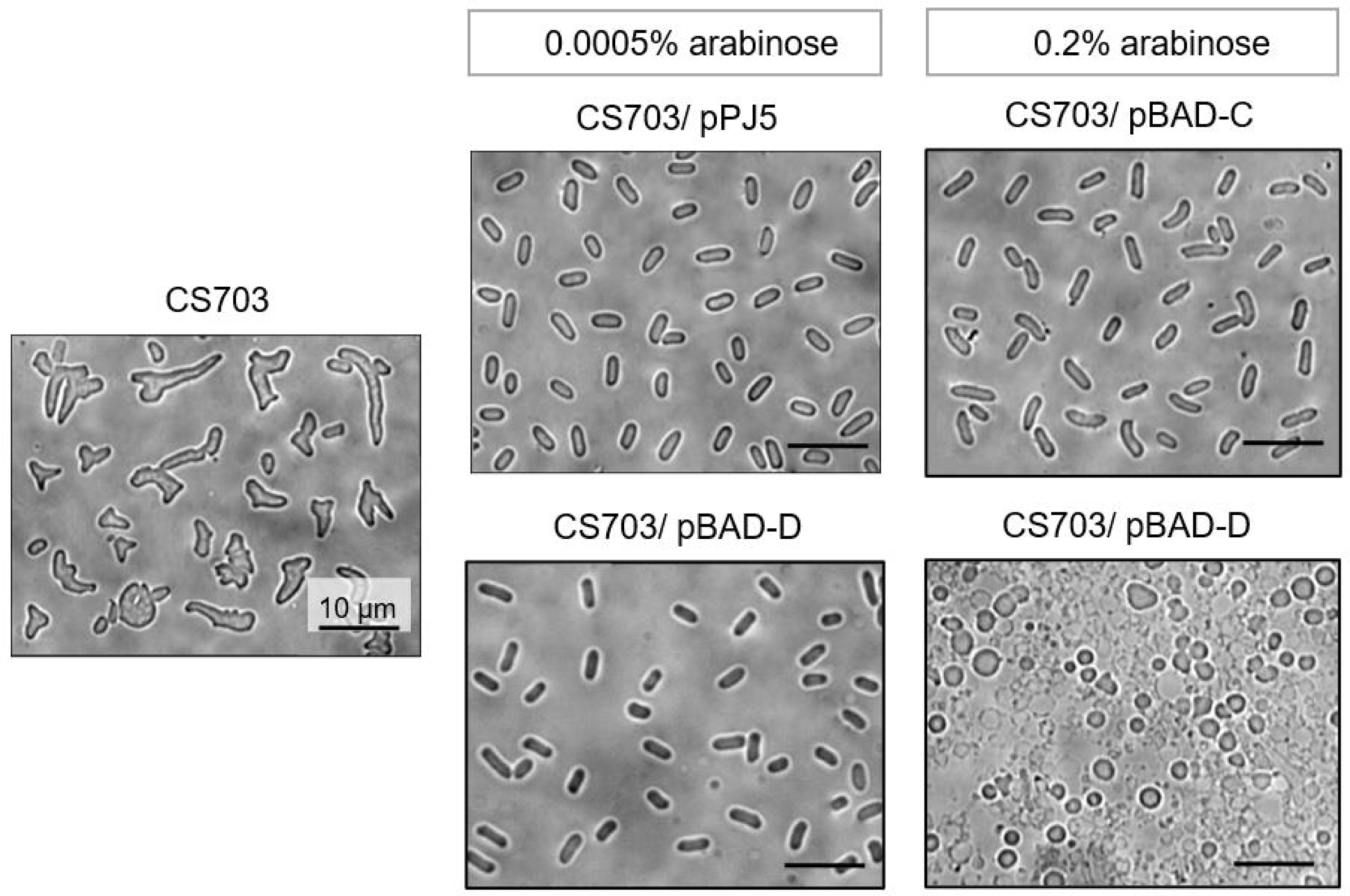
Ectopic expression of Ab PBP5/6 and Ab PBP6 can restore rod shape morphology in a septuple PBP mutant of *E. coli,* CS703. Bright field microscopic pictures (100x) of CS703, CS703/ pPJ5, CS703/ pBAD-C, CS703/ pBAD-D after inducing with different arabinose concentrations.

### Absence of dacC affects the growth rate of A. baumannii

To assess the effect of the absence of *dacC* and *dacD* on the fitness of *A. baumannii*, we monitored the growth of the deletion mutants. We observed that the growth rate was reduced in the cells devoid of *dacC* (Δ*dacC* and Δ*dacC*Δ*dacD* mutants) as compared to the parent (*A. baumannii* AB19606). However, the absence of *dacD* had no such effect (**Fig. S2**). The results indicate that although the viability of the cell is unaltered in the absence of *dacC* and *dacD*, *dacC* plays a vital role in maintaining the fitness of *A. baumannii*.

### Absence of AbDacC results in the loss of rod-shaped property

Unlike the wild type (WT) AB19606 cells, which were coccobacilli, the cells devoid of *AbdacC* lost the rod-shaped property, and the mutants appeared spherical (coccus like shape) (**Fig. 2**), which is corroborating with the previous findings (Geisinger et al., 2020). Interestingly, the loss of *AbdacD* did not alter the cell shape. The result indicates the possibility of the involvement of *AbdacC* in the process of maintaining a rod cell shape that might be the reason behind the spherical appearance in the Δ*dacC* deletion mutant, though further investigation may be required in future to critically justify the phenomenon. It is reported earlier that PBP deletions, especially, the LMM PBPs, might affect the biofilm forming ability of the cells (Gallant *et al*, 2005). Accordingly, we have attempted to investigate the influence of morphological changes in *A. baumannii* Δ*dacC* on its biofilm forming ability.

**Figure 2.**
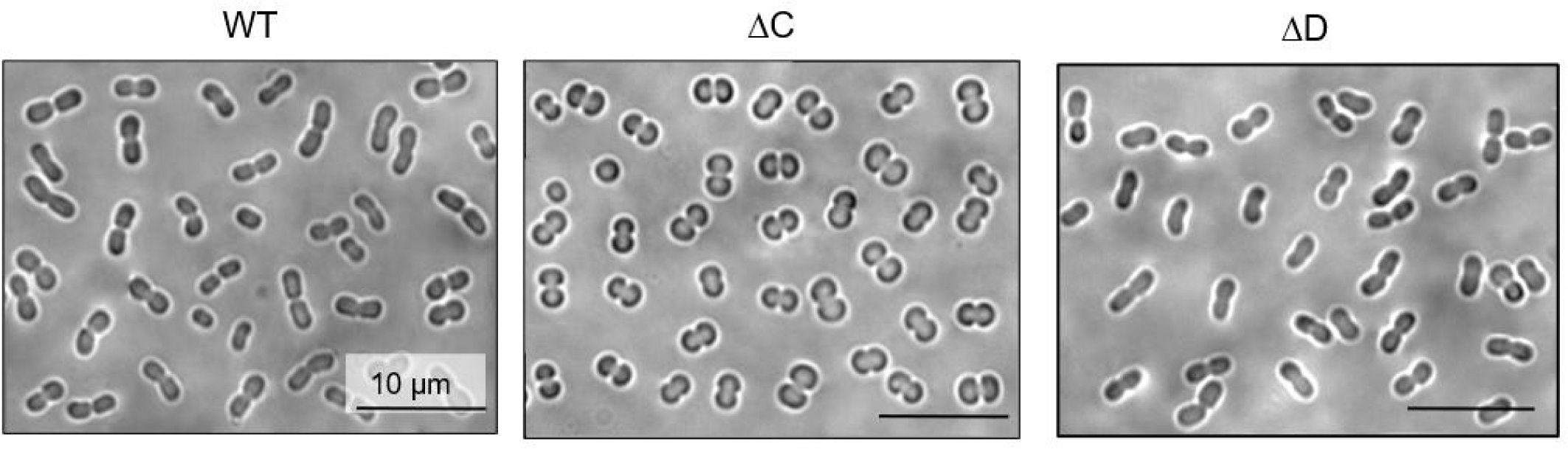
Microscopic analysis of the bacterial cell shape depicting the loss of rod-shaped property upon deletion of *dacC*. Bright Field microscopic pictures (100x) of *A. baumannii* AB19606 (WT), the DD-CPase knockout mutants AB19606Δ*dacC* (ΔC), AB19606Δ*dacD* (ΔD) and the complemented strain AB19606Δ*dacC*/pBAD-C (ΔC/ C). The lower panel of WT and ΔC are fluorescent images of the cells stained with FM4-64 (cell membrane) and DAPI (nuclear material).

### Biofilm production is reduced upon dacC deletion of A. baumannii

Biofilm formation is an important phenomenon in the development of *A. baumannii* infections as well as antimicrobial resistance. The alteration in morphology of the cell that is associated with DD-CPase activity can directly or indirectly influence the biofilm forming ability of the bacteria (Pandey et al., 2018b). In view of that, we investigated the roles of these PBPs on whether they can influence the biofilm formation of *A. baumannii*. We observed a reduction of biofilm formation by ∼35% in the absence of *dacC* (i.e., in Δ*dacC* mutant of AB19606) than that of wild-type. Trans-complementation of the mutants with the wild type *dacC* allele significantly restored the parental phenotype. However, deletion of *dacD* had only a slight effect on biofilm formation (**Fig. 3a, b**).

**Figure 3.**
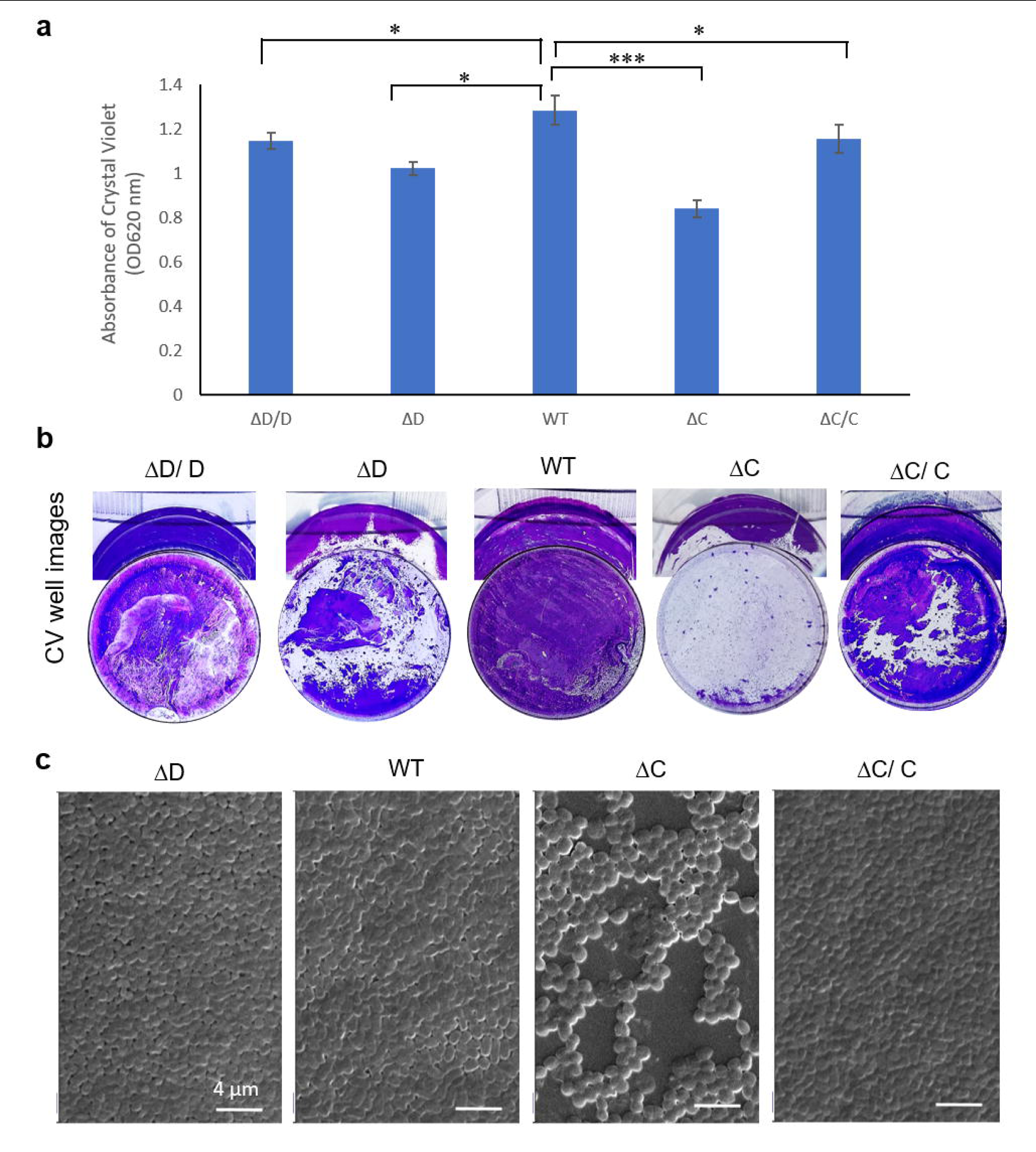
Absence of *dacC* in *A. baumannii* leads to reduced biofilm formation. (a) Quantitative analysis of biofilm formation by crystal violet (CV) staining of the strains *A. baumannii* AB19606 (WT) and the DD-CPase knockout mutants AB19606Δ*dacC* (ΔC) and AB19606Δ*dacD* (ΔD) and the complemented strains AB19606Δ*dacC*/pBAD-C (ΔC/ C) and AB19606Δ*dacD*/pBAD-D (ΔD/ D). Strains were grown in LB at 37 °C under static condition before staining with CV. Error bars are standard deviations of three replicates. A two-tailed, unpaired t test was used to determine the statistical relevance. *, P<0.05, ***, P<0.001 (b) Images of the CV-stained biofilms in the wells. (c) Scanning electron microscopic images of the biofilms. Scale bars represent 4 µm.

The quantitative data obtained from the *in vitro* biofilm assay using crystal violet (CV) further strengthened by the microscopic analysis of the biofilms. When the biofilms were observed under a scanning electron microscopy (SEM) (**Fig. 3c**), WT cells were found tightly packed and forming dense biofilm. On the other hand, the inter-cellular spacing was more in the low biofilm forming *dacC* mutant. Therefore, it can be inferred that the deletion of *dacC* significantly lowers the biofilm formation in AB19606.

### AB_DacC (PBP5/6) imparts intrinsic β-lactam resistance to *A. baumannii* whereas AB_DacD (PBP6b) does not affect β-lactam susceptibility

*E. coli* cells become sensitive to β-lactams in the absence of PBP5 (*dacA*), and ectopic complementation of PBP5 reverses the lost resistance (Sarkar et al., 2010). In view of that, to investigate whether the DD-CPases of *A. baumannii* also had a similar role, both *dacC* and *dacD* were expressed *in trans* in AM15 (*E. coli*ΔPBP5) cells. AM15 is approx. 2 to 8-fold more sensitive towards β-lactam agents than that of the parent *E. coli* CS109 due to absence of an intrinsic beta-lactam resistance factor, *ampC* (Sarkar et al., 2010). *In trans* expression of *E. coli dacA* (PBP5) complements the loss to some extent. Likewise, expression of *A. baumannii dacC* reversed the lost β-lactam resistance in AM15, indicating its ability in maintaining the intrinsic β-lactam resistance of *E. coli* (**Table S3**). However, expression of *dacD* was unable to complement the lost resistance.

Since, *A. baumannii dacC* could restore the lost β-lactam resistance of *E. coli*, it was imperative to study the effect of absence of *dacC* in *A. baumannii* on its β-lactam susceptibility. To investigate this, we estimated the antibiotic sensitivities of the mutants and complemented strains by determining MICs. Deletion of *dacC* results in the sensitization *of A. baumannii* AB19606 (**Table 2**) towards β-lactam agents by 2 to 8-fold as compared to their wild type counterparts. Ectopic expression of *dacC* increases the resistance and restores it to wild type levels. However, *in trans* expression of *dacD* did not exert a similar effect. Overall, the results indicate that unlike PBP6b *(dacD)*, the PBP5/6 (*dacC*) imparts intrinsic β-lactam resistance to *A. baumannii*.

**Table 2.** β-lactam susceptibility (MIC µg mL^-1^) of *A. baumannii* DD-CPase mutants.

|  | Penicillin |  |  |  |  | Carbapenem |  | Cephalosporin |  |  |  |  |
| --- | --- | --- | --- | --- | --- | --- | --- | --- | --- | --- | --- | --- |
| Strains | PENG | OXA | AMX | TIC | PIP | DOR | MER | CXM | FOX | CAZ | CFS | CTX |
| WT (19606) | >2000 | 500 | 1000 | 16 | 500 | 1.6 | 1.6 | 500 | 500 | 16 | 250 | 64 |
| $\Delta\text{C}$ | 2000 | 250 | 500 | 8 | 64 | 0.8 | 0.4 | 250 | 250 | 8 | 64 | 32 |
| $\Delta\text{C}/\text{C}$ | >2000 | 500 | 1000 | 16 | 500 | 1.6 | 1.6 | 500 | 500 | 16 | 250 | 64 |
| $\Delta\text{C}/\text{D}$ | 2000 | 250 | 500 | 8 | 64 | 0.8 | 0.4 | 250 | 250 | 8 | 64 | 32 |
| $\Delta\text{D}$ | >2000 | 500 | 1000 | 16 | 500 | 1.6 | 1.6 | 500 | 500 | 16 | 250 | 64 |
| $\Delta\text{D}/\text{D}$ | >2000 | 500 | 1000 | 16 | 500 | 1.6 | 1.6 | 500 | 500 | 16 | 250 | 64 |
| $\Delta\text{C}\Delta\text{D}$ | 2000 | 250 | 500 | 8 | 64 | 0.8 | 0.4 | 250 | 250 | 8 | 64 | 32 |
| $\Delta\text{C}\Delta\text{D}/\text{C}$ | >2000 | 500 | 1000 | 16 | 64 | 1.6 | 1.6 | 500 | 500 | 16 | 250 | 64 |
| $\Delta\text{C}\Delta\text{D}/\text{D}$ | 2000 | 250 | 500 | 8 | 64 | 0.8 | 0.4 | 250 | 250 | 8 | 64 | 32 |

[PENG: Penicillin G; OXA: Oxacillin; AMX: Amoxicillin; TIC: Ticarcillin; PIP: Piperacillin; DOR: Doripenem; MER: Meropenem; CXM: Cefuroxime; FOX: Cefoxitin; CAZ: Ceftazidime; CFS: Cefsulodin; CTX: Cefotaxime]

### dacC and dacD are expressed in different growth phases

The redundancy of DD-Carboxypeptidases can be related to their need at different growth phases of bacteria (Buchanan and Sowell, 1982b; Peters et al., 2016a). Therefore, the expressions of *dacC* and *dacD* in the strain AB19606 were assessed in different growth phases. The 16s rRNA was used as the endogenous control. The RNA expression studies revealed that the *dacC* expression started from early logarithmic phase and the maximum expression was at the mid to late logarithmic phases, which started declining at the early stationary phase. On the other hand, *dacD* expression started nearly at the late logarithmic phase and reached the peak at the late stationary phases (**Fig. 4**). Therefore, it can be summarized as *dacC* is predominantly expressed in the exponential phase whereas *dacD* is expressed in the stationary phase.

**Figure 4.**
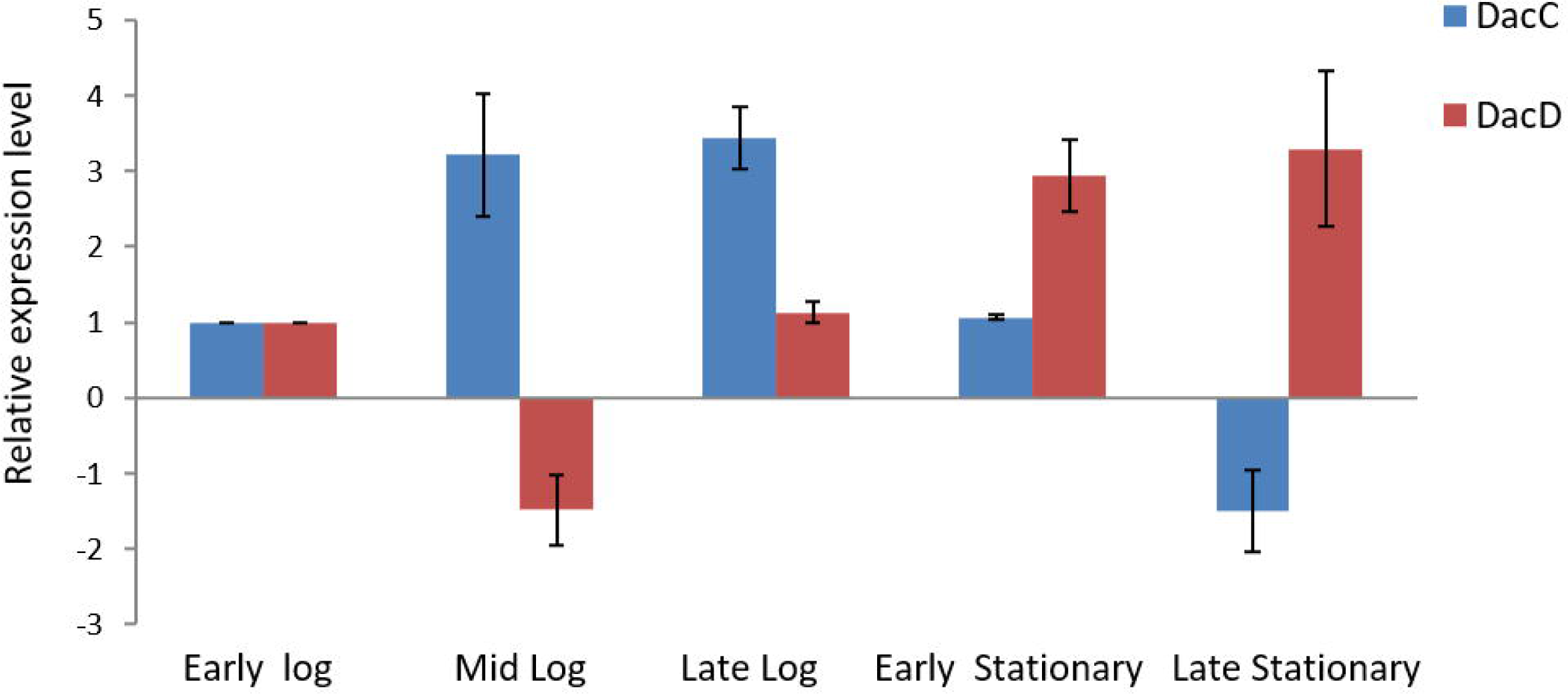
qPCR-based transcript analysis depicting differential expression of *dacC* and *dacD* in different growth phases of *A. baumannii* AB19606. The endogenous control used was 16S rRNA.

### The in vitro DD-CPase activity of DacD (PBP6b) is higher than DacC (PBP5/6) in A. baumannii

PBPs are ectoproteins that localized in the periplasmic space and are anchored to the inner membrane of the bacterial cell wall. Therefore, the signal peptides and the membrane anchors had to be removed to solubilise the proteins. The soluble counterparts of the proteins PBP5/6 (sDacC) and PBP6b (sDacD) containing 348 aa and 388 aa, respectively, were purified through affinity chromatography (see methods). The molecular mass of sPBP5/6 was 40.9 kDa and that of sPBP6b was 45.5 kDa as determined through MALDI analysis **(Fig. S3)**. Circular dichroism (CD) data showed that the proteins were in the native conformation and the percentages of α-helix, β-sheet, turn and random coil were 27.5%, 13.3%, 32.4% and 26.8%, respectively, for sDacC and 24.3%, 8.7%, 30.6% and 36.4% for sDacD (**Table S4**).

The DD-CPase activity was determined by using a peptidoglycan mimetic pentapeptide substrate, L-Ala-c-D-Glu-L-Lys-D-Ala-D-Ala (Chowdhury et al., 2010). During the DD-CPase assay the different concentrations of pentapeptide substrate was hydrolysed by the PBP and the concomitant release of the terminal D-Ala was measured spectrophotometrically. Interestingly, though sDacC has ∼3 times higher affinity towards the pentapeptide substrate as compared to sDacD, the catalytic efficiency (*k*_cat_/*K*_m_) of sDacD (∼6.3 s^-1^ mM^-1^) was nearly 1.75 times than that of sDacC (∼3.6 s^-1^ mM^-1^) (**Table 3**). The results implied that sDacD has higher *in vitro* DD-CPase activity than that of sDacC for the peptidomimetic substrate (**Fig. 5**). These results substantiated our earlier *in vivo* studies, in case of *dacD,* where higher inducer concentration led to lysis of the host *E. coli* cells.

**Figure 5.**
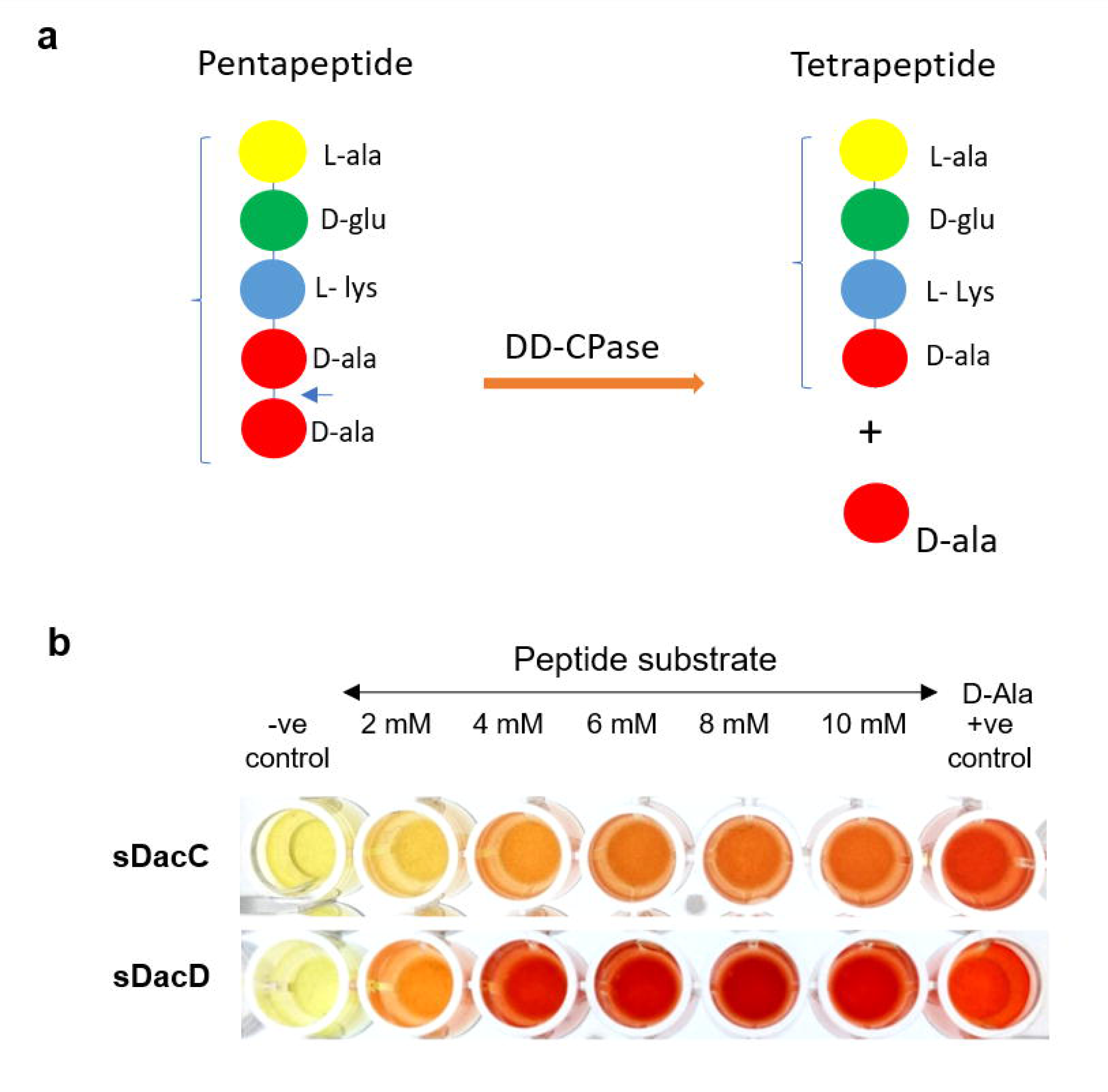
*in vitro* DD-CPase Assay. a. Schematic representing the enzymatic conversion of the pentapeptide-mimetic substrate to tetrapeptide and D-Alanine in the *in vitro* DD-CPase assay. b. DD-CPase assay using a peptidoglycan-mimetic pentapeptide substrate depicting enhanced DD-CPase activity of sDacD.

**Table 3.** Kinetic parameters of the DD-CPase activity of the purified PBPs with pentapeptide substrate.

| Protein | (Pentapeptide)<br>L-Ala-γ-D-Glu-L-Lys-D-Ala-D-Ala | | |
| --- | --- | --- | --- |
| | $K_m$<br>(mM) | $k_{cat}$<br>(s<sup>-1</sup>) | $k_{cat}/K_m$<br>(s<sup>-1</sup> mM<sup>-1</sup>) |
| **sDacC** | 0.32 ± 0.12 | 1.16 ± 0.14 | 3.63 |
| **sDacD** | 1.09 ± 0.17 | 6.9 ± 0.8 | 6.33 |

### Soluble DacC (sPBP5/6) shows a high acylation rate as well as a significantly high deacylation rate towards penicillin

Further to explore the possible mechanism of sDacC in imparting intrinsic beta-lactam resistance, we investigated its role in acylation and deacylation of beta-lactam substrates. β-lactams are substrate analogues of the D-Ala-D-Ala terminus of the peptide side chain in PG, and bind to PBPs. The efficiency with which penicillin binds to the PBP, to form the acyl-enzyme intermediate, is determined by the acylation rate. Here, we used the fluorescent penicillin, Bocillin FL to determine the acylation rates of sDacC, i.e., sPBP5/6. The acylation rate constant (*k_2_/K*) of sDacC was 2800 ± 190 M^-1^s^-1^. During the deacylation reaction, the covalently bound acyl-enzyme complex is hydrolysed and the inactive β-lactam is released. The deacylation rate is estimated by measuring the first-order rate constant *k_+3_*. The deacylation rate of sDacC was (260 ± 45) x 10^-5^ s^-1^. Moreover, we observed that unlike sDacD (sPBP6b), when sDacC was incubated with Bocillin for longer durations, the intensity of the Bocillin-labelled bands were diminished, implicating that the bound Bocillin was being hydrolysed by sDacC manifesting very high deacylation efficiency of sDacC. The higher deacylation rate indicates the efficiency of beta-lactam hydrolysing by sDacC. Meanwhile, some DD-CPases are also reported to possess β-lactamase activity that makes them dual enzymes (Bansal et al., 2015). However, such β-lactamase activities are usually observed against different class of β-lactam. To check whether sDacC has β-lactamase activities against different classes of β-lactams, we assessed the hydrolysing capacity of sDacC against penicillins, cephalosporins and carbapenems. Herein, we observed that sDacC hydrolysed all of these antibiotics classes, which is in synchrony with the antibiotic sensitivity analysis data. The kinetic parameters depicting the β-lactam hydrolyzing efficiency of sDacC are listed in **Table 4**. *In vitro* kinetic assays further validated the fact that sDacC has β-lactamase activity. Therefore, it can be concluded that DacC (PBP5/6) of *A. baumannii* is a dual-function enzyme manifesting β-lactamase and DD-carboxypeptidase activities.

**Table 4.**
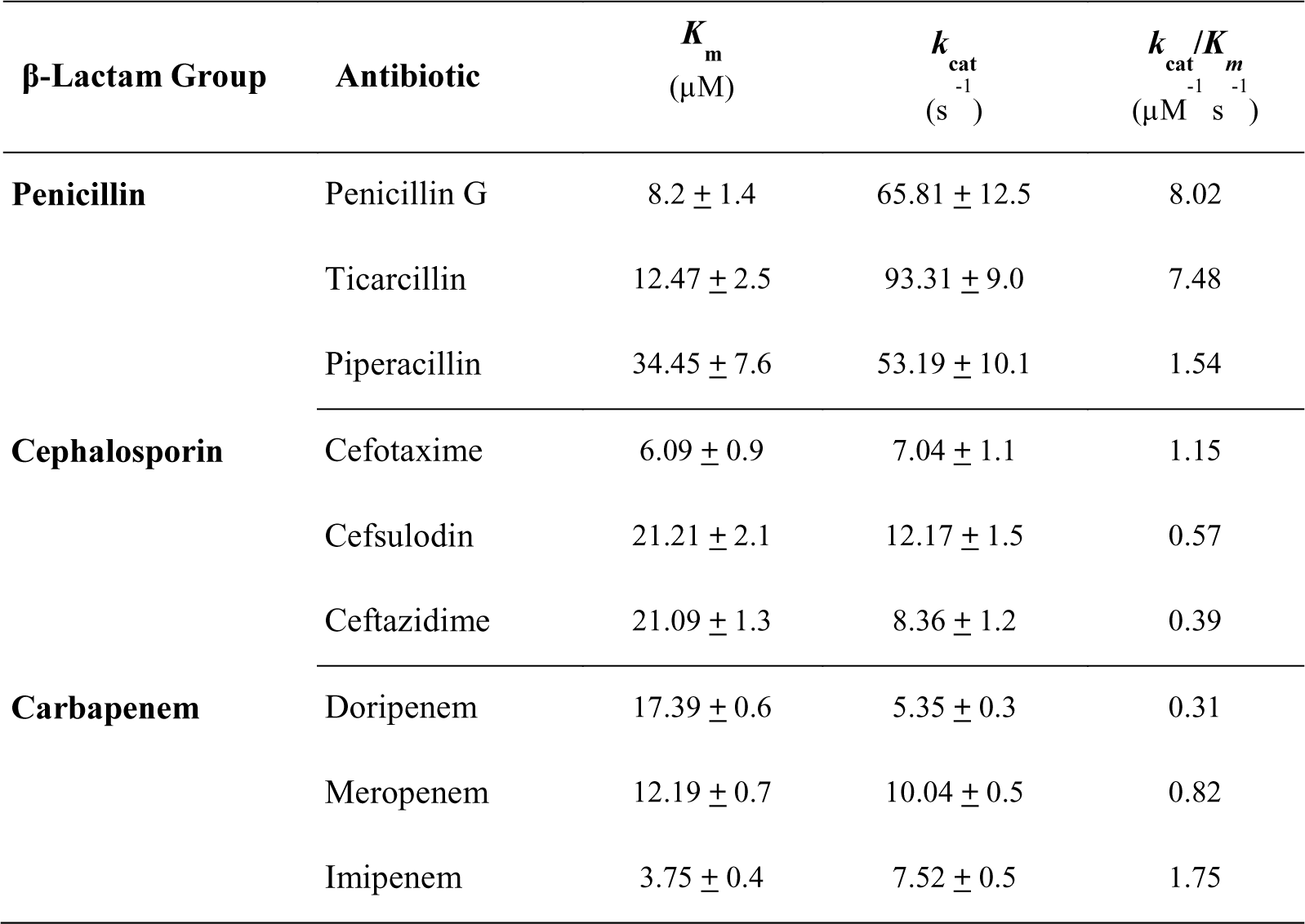
Kinetic parameters for the β-lactamase activity of purified sDacC.

| β-Lactam Group | Antibiotic | $K_m$<br>($\mu\text{M}$) | $k_{cat}$<br>($\text{s}^{-1}$) | $k_{cat}/K_m$<br>($\mu\text{M}^{-1}\text{s}^{-1}$) |
| --- | --- | --- | --- | --- |
| Penicillin | Penicillin G | $8.2 \pm 1.4$ | $65.81 \pm 12.5$ | 8.02 |
| | Ticarcillin | $12.47 \pm 2.5$ | $93.31 \pm 9.0$ | 7.48 |
| | Piperacillin | $34.45 \pm 7.6$ | $53.19 \pm 10.1$ | 1.54 |
| Cephalosporin | Cefotaxime | $6.09 \pm 0.9$ | $7.04 \pm 1.1$ | 1.15 |
| | Cefsulodin | $21.21 \pm 2.1$ | $12.17 \pm 1.5$ | 0.57 |
| | Ceftazidime | $21.09 \pm 1.3$ | $8.36 \pm 1.2$ | 0.39 |
| Carbapenem | Doripenem | $17.39 \pm 0.6$ | $5.35 \pm 0.3$ | 0.31 |
| | Meropenem | $12.19 \pm 0.7$ | $10.04 \pm 0.5$ | 0.82 |
| | Imipenem | $3.75 \pm 0.4$ | $7.52 \pm 0.5$ | 1.75 |

### Differential affinity of purified AbDacC and AbDacD for β-lactams and β-lactamase inhibitors

PBPs are the targets of β-lactam antibiotics, though they have difference in affinities for different β-lactams and the binding affinities of PBPs vary from species to species. *A. baumannii* has emerged as a nosocomial pathogen but the binding affinity of β-lactams with AB PBPs is still not explored extensively. The affinities of different β-lactams and β-lactamase inhibitors for sDacC and sDacD were determined experimentally by monitoring the inhibition of PBP acylation by Bocillin FL.

The binding of the drugs to the PBPs was quantified by adding increasing concentrations of the antibiotics/inhibitors to the purified soluble proteins prior to addition of Bocillin. The fluorescence intensities of the Bocillin labelled PBPs was quantified to estimate the relative binding affinity (50% inhibitory concentration, IC_50_). The IC_50_ values for different β-lactams and inhibitors for sDacC (sPBP5/6) and sDacD (sPBP6b) are presented in **Table 5**.

**Table 5.** Relative binding (IC_50_) of β-lactams and β-lactamase inhibitors to purified DacC and DacD (µM)

| Category | Antibiotic | sDacC | sDacD |
| --- | --- | --- | --- |
| Penicillins | Oxacillin | 95.9 | 417.1 |
|  | Piperacillin | 539.2 | 3280.8 |
| Cephalosporins | Cefuroxime | 12.9 | 160.8 |
|  | Cefexime | 20.9 | 52.8 |
|  | Cefsulodin | 6.3 | 8.6 |
| Carbapenems | Doripenem | 9.2 | 1.7 |
|  | Meropenem | 15.7 | 2.7 |
| β-lactamase Inhibitors | Clavulanic acid | 585.6 | 5.8 |
|  | Tazobactam | 1828.2 | 1591.5 |
|  | Sulbactam | 183.8 | 3.8 |

The selectivity profiles of both the PBPs differed substantially. The acylation ability of carbapenems (doripenem and meropenem) was good for both the DD-CPases, though sDacD was inhibited more efficiently. Representative gel band quantitation graphs are depicted in **Fig. S4a**. The β-lactamase inhibitors (clavulanic acid, sulbactam) demonstrated selective inhibition of sDacD as the IC_50_ values were significantly lower for sDacD as compared to that of sDacC. Both sPBP5/6 and sPBP6b showed high affinity for cephalosporins. The affinity of sPBP5/6 was depicted relatively higher for cefuroxime and cefixime than that of sPBP6 (**Fig. S4b**). However, both sDacC (sPBP5/6) and sDacD (sPBP6b) had high affinity for cefsulodin. The affinity of sDacC towards penicillins was relatively less as compared to its affinities for cephalosporins and carbapenems. Among the antibiotics tested, sDacD had the highest affinity for doripenem, followed by meropenem, sulbactam, clavulanic acid, and cefsulodin whereas the binding efficiency of sDacC was highest towards cefsulodin and doripenem followed by cefuroxime, meropenem, and cefixime.

## Discussion

The physiology of LMM-PBPs is remaining as a mystery still today, though various investigations were carried out in different strains. To study the physiological roles of putative DD-CPases, PBP5/6 (*dacC*) and PBP6b (*dacD)* of *A. baumannii*, we have created single and double deletion mutants of these genes. The loss of *dacC* and *dacD,* either individually or in combination, were not lethal for *A. baumannii*. These observations are in line with the previous reports on LMM PBPs indicating their non-essential nature for survival (Ghosh et al., 2008). Previously, it has been reported that the loss of single LMM-PBPs does not affect the viability of *Streptococcus pneumoniae (*Schuster *et al., 1990)* and *S. aureus* (PBP4)(Kozarich and Strominger, 1978). Also, *E. coli*, *N. gonorrhoeae*, and *B. subtilis* can tolerate the loss of all LMM PBPs (Popham et al., 1999; Stefanova et al., 2003). A possible reason for the non-essential nature of LMM PBPs could be the genetic multiplicity of the genes encoding them, as well as their abundance, which facilitates the substitution of the loss of one LMM PBP by another. However, there are exceptions, as recently Ealand *et al*. reported the essential nature of an LMM PBP in *M. smegmatis.* Attempts to delete the *dacB* (MSMEG_6113) gene, which encodes an LMM-PBP from *M. smegmatis* are unsuccessful which indicates its essential role in cell viability (Ealand et al., 2019).

Though not lethal to *A. baumannii*, the absence of *dacC* affected the growth rate but not *dacD.* The growth of *E. coli* is unaltered even in the absence of all its LMM PBPs (Denome et al., 1999). Similarly, *B. subtilis* cells devoid of nearly all the DD-CPases grow normally (Popham et al., 1999). However, *N. gonorrhoeae* cells lacking both the DD-CPases PBP3 and PBP4 grow slower but the growth is unaltered in the absence of single DD-CPases (Stefanova et al., 2003)). Similarly, *V. cholerae* devoid of the PBP5/6 homologue DacA-1 displays slow growth, though the lack of *dacA-2*, *dacB*, and *pbpG*, do not affect the growth rate (Möll et al., 2015). *M. smegmatis* strains lacking one DD-CPase gene (the single knockouts of MSMEG_2432, MSMEG_2433 and MSMEG_1661) display no growth rate defects (Pandey et al., 2018a; Ealand et al., 2019). However, the absence of both the DD-CPases MSMEG_2432 and MSMEG_2433 results in a 16% lower growth rate in *M. smegmatis* (Pandey et al., 2018a). The absence of PBP5/6 in *L. monocytogenesb* (Korsak et al., 2005) and PBP3 in *S. pneumoniae* (Korsak et al., 2010) reduces the growth rate. Therefore, it can be speculated that the role of an LMM PBP in the bacterial fitness and growth is species specific.

### Loss of a single non-essential PBP can also significantly impact the cell morphology

The cell shape studies reveal that the absence of *dacC* results in the loss of the rod-shaped property in *A. baumannii,* which is in line with the previous studies (Geisinger et al., 2020). The otherwise coccobacillus AB19606 cells (mostly occurring in diplo coccobacillus form) appear spherical (diplococcus) when *dacC* is deleted. There are several reports which depict the importance of DD-CPases in maintaining normal cell morphology. The absence of a single LMM PBP does not alter the cell shape of *E. coli* and the cells devoid of PBP5 have normal cell shape (Nelson and Young, 2001a), though the cells display aberrant morphology upon deletion of any two other LMM PBP genes along with PBP5. The mutant cells have altered cell diameter with kinks, bends, and branches (Potluri et al., 2010).

The absence of a DD-CPase affecting the cell shape is also reported. *N. gonorrhoeae* cells lacking both PBP3 (similar to *E. coli* PBP4) and PBP4 are morphologically different from the parent and the single mutants. The double knockout comprises both large and small cells implicating defects in cell division (Stefanova et al., 2003). The morphology of *V. cholerae* is altered in the cells lacking *dacA-1,* unlike the other LMM PBPs. The cells of the Δ*dacA-1* strain are elongated and enlarged, and form aberrant poles and branches. As growth progresses, the defects become pronounced. The effect is reversed upon complementation with both DacA-1 and DacA-2 (Möll et al., 2015). Morphological changes like unusual septum formation and cell wall thickening are observed in *Streptococcus pneumoniae* PBP3 (similar to the *E. coli* PBP5) mutants and the cells grow in clumps as enlarged spheres (Schuster et al., 1990).

The PG layer or the murein sacculus determines the bacterial cell shape, which in turn is dependent on PBPs that catalyse the insertion of murein precursors into the wall. DD-CPases control the amount of pentapeptide substrate available to the PG synthetic transpeptidases (Peters et al., 2016b). Absence of DD-CPases leave a greater proportion of pentapeptide side chains in mature peptidoglycan, thereby altering the ratio of pentapeptide and tetrapeptide substrates that eventually affects PG synthesis. This could affect the competition between the elongation and septation pathways and the resulting imbalance or misplacement of peptidoglycan synthesis that might create the misshapen cells. Although the exact mechanism is unknown, the elevated pentapeptide levels in DD-CPase mutants might alter the geometry of the septal ring which might affect the localization and proper functioning of PG synthases which eventually affects the morphology of the bacteria. Additionally, it was also observed that the co-localization of HMM PBPs and FtsZ and FtsW was affected in the *S. pneumoniae* mutant lacking the DD-CPase PBP3 (Morlot et al., 2004). Our results further establish the vital role of DD-CPases in cell shape maintenance. Here, we observe that the cells become spherical upon the loss of *dacC* and the effect is reversed upon *in trans* complementation with *dacC*. Similar observations were made by Nelson *et al*. who reported that the ectopic expression of PBP5 in *E. coli* nullifies the cell shape abnormalities in *E. coli* cells devoid of LMM PBPs but the complementation with the other *E. coli* DD-CPase genes encoding PBP4, PBP6, or DacD are unable to restore normal cell shape (Nelson and Young, 2001a).

Biofilm production was also reduced in the mutant lacking *dacC*. Previously, a reduction in biofilm formation is reported in the double knockout *M. smegmatis* strain lacking MSMEG_2433 and MSMEG_2432 (Pandey et al., 2018a), though the biofilm production remains unaltered in the single DD-CPase mutant of *M. smegmatis* (Ealand et al., 2019). As the structural integrity of the cell depends on the PG layer, the altered PG structure in the absence of DD-CPase may possibly alter the surface topology and the arrangement of adhesin proteins that eventually affects the biofilm formation.

### DacC imparts intrinsic β-lactam resistance to A. baumannii

Although alterations in HMM-PBPs are responsible for clinically relevant β-lactam resistance, LMM-PBPs also play a role in imparting intrinsic resistance to some antibiotics (Sarkar et al., 2010; Sarkar et al., 2011). Here, we observe that *dacC* deletion mutants become sensitive to β-lactams. There are reports that indicate the role of DD-CPases in intrinsic β-lactam resistance (Sarkar et al., 2010). In *E. coli* among all the DD-CPases, only PBP5 plays a role in imparting intrinsic β-lactam resistance, as the deletion of PBP5 sensitizes *E. coli* cells to β-lactams by four to eightfold and the effect is restored upon complementation with PBP5 (Sarkar et al., 2010). The ectopic expression of PBP6 is unable to restore the lost β-lactam resistance, though overexpression of DacD partially restores the lost resistance (Sarkar et al., 2011). Similarly, the DD-CPase PBP5 of *P. aeruginosa* possesses β-lactamase activity (Smith et al., 2013). The inactivation of PBP4 (*dacB*) enhances the β-lactam resistance of *P. aeruginosa* by inducing AmpC expression (Moya et al., 2009). All these reports have strengthened the fact that some LMM PBPs impart intrinsic resistance to β-lactams and in *A. baumannii*, the LMM PBP that has such property is the DacC. Therefore, it is evident from the obtained results that DacC might play an additional role in quenching β-lactam activity, most likely by binding to these antibiotics efficiently to mask their effect on the essential PBPs (Sarkar et al., 2010) and/or by hydrolysing β-lactams (probably partially) to minimize their availability within the periplasm that in turn might protect the essential PBPs from binding to these β-lactams.

### Differential expression of DacC and DacD at different growth phases justifies the significant role of these PBPs in maintaining the physiology of A. baumannii

The RNA expression studies revealed that in *A. baumannii, dacC* is expressed in both exponential and early stationary phases, whereas *dacD* is expressed predominantly in the stationary phase (**Fig. 4**). It is observed that bacterial species that harbour multiple DD-CPases express them at different phases or conditions, which explains the reason behind their genetic multiplicity. *E. coli* encodes four PBPs with DD-CPase activity, among which the expression of PBP5 occurs in the exponential phase (Santos et al., 2002) whereas PBP6b expresses in the mid-logarithmic phase of bacterial growth (Baquero et al., 1996) and PBP6 during the stationary phase (Buchanan and Sowell, 1982a; Van der Linden et al., 1992). Similarly, the expression of the PBP5 homologue *dacA*-1 in *V. cholerae* is ∼3× higher during the exponential phase (Möll et al., 2015). *B. subtilis* cells lacking PBP5 are normal in the exponential phase but become shorter in the stationary-phase of growth and produce spores with an altered cortex structure (Todd et al., 1986). Therefore, the physiological impact of a particular PBP depends on the growth phase in which it is expressed. PBPs those express in the exponential phase play crucial roles in cellular physiology and could act as a potential drug target for actively growing cells, whereas the ones that are expressed in later stages can serve as targets for eradicating dormant persister cells in biofilms.

### *In vitro* studies corroborated the *in vivo* observations

The *in vivo* studies revealed that *A. baumannii* DacC and DacD are DD-CPases. To further validate the results, the proteins were purified and *in vitro* kinetics is assessed to evaluate their catalytic activities. Therefore, the catalytic activity of the DD-CPases is determined by assessing the hydrolysis of the peptidoglycan-mimetic pentapeptide substrate and purified sDacD exhibits strong DD-CPase activity. However, the DD-CPase activity of sDacC is moderate. The results confirm that these two putative DD-CPases have DD-CPase activity and explain the differential cell shape maintenance properties of both the PBPs. In *E. coli*, the *in vitro* DD-CPase activity of sDacD (*k*_cat_∼ 0.13 s^-1^) is less as compared to *E. coli* sPBP5 (*k*_cat_∼ 1.4 s^-1^) (Chowdhury*, et al.*, 2012). The fact that DacD is the stronger DD-CPase in *A. baumannii* and is a weaker DD-CPase in *E. coli* indicates that similar proteins might function differently in different species. Further, *in silico* analysis helps us to understand the molecular basis of this differential activity of DD-CPases observed in both the proteins.

To explore the binding interaction of the pentapeptide substrate with the protein, docked complexes of both DacC and DacD with pentapeptides are analysed (See Supplementary Materials for molecular modelling & bioinformatic analysis methodology). The docking complex of DacD (**Fig. 6a**) revealed that the pentapeptide binds precisely in the active site pocket of the enzyme within a close proximity of the catalytic residues. Formation of a strong interacting network is seen due to the presence of strong hydrogen bond bridges with the catalytic residues of the protein (indicated by dotted lines) in **Fig. 6b**. A very stable intermolecular environment is formed between the protein and substrate through hydrogen bond formation with several residues, namely, Lys 63, Asn 124, Ala 123, Glu 98, Asp 97, Trp 96, Ser 99, thereby facilitating the DD-CPase activity. On the contrary, a significantly small number of interacting bonds was found in the DacC-pentapeptide docked complex, implicating a weak protein-substrate interaction which corroborates with the *in vivo* data. Additionally, the orientation of the pentapeptide substrate within the active site also affects the activity (Chowdhury et al., 2011). Further analysis of the protein-ligand interactions is done using Lig-plot. The Lig-plot analysis of the proteins indicated that the surface interaction of DacD is higher than that of DacC (**Fig 6c**). Although both the proteins have affinity towards the pentapeptide substrate, the number of hydrophobic interactions is more in case of DacD as compared to DacC, further justifying the reason behind the strong DD-CPase activity of DacD.

**Figure 6.**
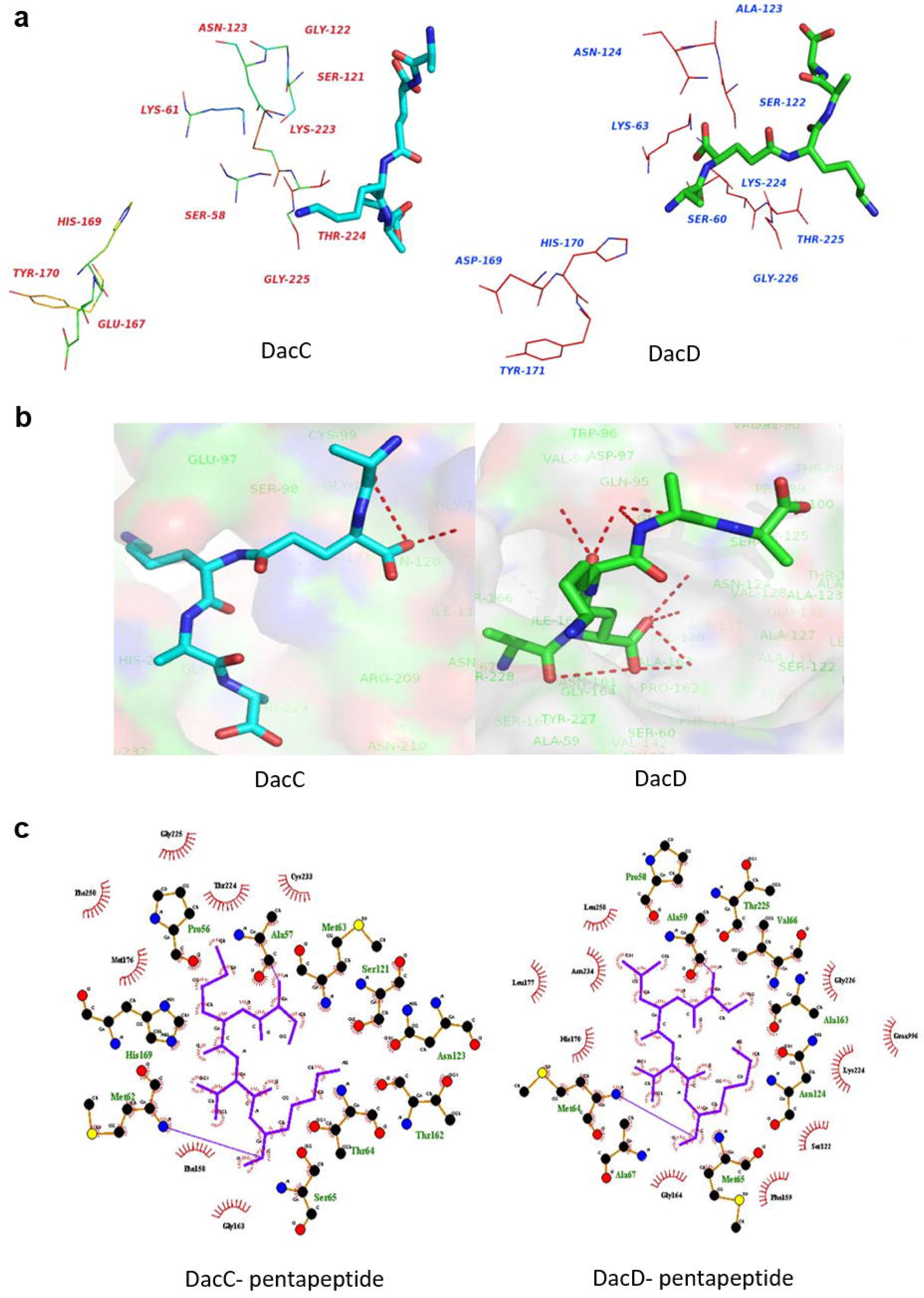
Docking results of interaction complexes of DacC and DacD with the pentapeptide substrates. **a.** The arrangements of active site residues around the pentapeptide substrate and conformations of the pentapeptide substate with DacC and DacD after docking analysis. **b.** Hydrogen bond interactions between the substrate and the proteins DacC and DacD. **c.** Lig-plot analysis for the interactions of the pentapeptide substrates with selected residues of DacC and DacD.

The rate of acyl-enzyme complex formation determines the efficiency with which a PBP binds to a β-lactam. The ability to bind to penicillin was estimated by determining the acylation rate with the substrate (Bocillin). We observed that the acylation rate of *A. baumannii* sDacC was ∼2800 M^-1^ s^-1^. Previously, it was seen that the acylation rates of *E. coli* sPBP6 towards Bocillin was ∼1900 M^-1^ s^-1^ whereas that of sPBP5 was 800 M^-1^ s^-1^and that of sDacD was 450 M^-1^ s^-1^(Chowdhury et al., 2012a and Chowdhury et al., 2012b). This indicates that the acylation rate of sDacC of *A. baumannii* is significantly high for penicillin.

During deacylation, the β-lactam-PBP complex breaks down, with the release of the cleaved antibiotic and the free enzyme. Proteins that have higher rates of deacylation hydrolyse the drugs faster and render the enzyme free. The deacylation rate of sDacC was very high which indicated its penicillin hydrolytic activity. In *E. coli,* the ability to deacylate Bocillin was highest for sPBP5, followed by sDacD and lastly sPBP6 (Chowdhury et al., 2012a). The PBP MSMEG_2433 in *M. smegmatis* which has β-lactamase activity also displays a high deacylation rate with Bocillin (Bansal et al., 2015).

The β-lactamase like property of sDacC is further confirmed through the β-lactam hydrolysis assay. It is found capable of hydrolysing penicillins, cephalosporins and carbapenems. *E. coli* PBP5 does not possess β-lactamase activity, under physiological conditions. However, *P. aeruginosa* PBP5 has been reported to possess a broad spectrum β-lactamase activity and can hydrolyse penicillins, cephalosporins, and carbapenems (Smith et al., 2013). MSMEG_2433 which is a DD-CPase in *M. smegmatis* is also reported to hydrolyse oxyimino-cephalosporins, aztreonam and penicillins, though it does not hydrolyse imipenem (Bansal et al., 2015). These results indicate that *A. baumannii* DacC is a dual enzyme and possesses both β-lactamase and DD-CPase activities. However, DacD might only be a DD-CPase.

PBPs and β-lactamases share both structural and sequence similarities and are believed to have a common ancestral origin (Ghuysen, 1991). They share common conserved motifs, protein folds, sequence similarities along with the catalytic mechanism which involves an active site serine residue (Meroueh et al., 2003). The conserved functional motifs of PBPs (also present in β-lactamases), which include SXXK tetrad, SXN triad and a KTG triad (Ghuysen, 1991) are identified in both DacC and DacD. The amino acid sequence similarity between *A. baumannii* DacC and DacD is 31.2%. To further identify the structural similarities/differences between DacC and DacD, the available 3D structures of both the proteins were superimposed and the RMSD (root mean square deviation) value was calculated. The difference between the RMSD score, predicted by the TM-align software (Zhang and Skolnick, 2005), for the tertiary structures of both the proteins is found to be 1.69, which strongly supports that there is a significant level of structural difference between the two predicted 3D protein structures.

Moreover, the active site groove volumes of DacC and DacD were predicted using the surface topology analysis server CASTp. The groove volume of DacC is estimated to be 610.6 Å^3^, whereas for DacD it is 1107.8 Å^3^. PBPs and beta-lactamases share a common penicilloyl serine transferase fold. In case of PBPs, the active site scoop is situated at the bottom of the elongated cleft and wider in nature whereas in β-lactamases it is narrower and within a surface pocket. This difference in the active site clearly indicates that PBPs have to strictly accommodate two stem peptides within their catalytic groove volume whereas β-lactamases have to accommodate just the β-lactam antibiotics (Macheboeuf et al., 2006). The limited spatial arrangement of the binding pocket possibly provides inadequate space for the proper placing of the pentapeptide substrates. This major difference in the active site pocket suggests that DacD has an elongated active site cleft supporting its DD-CPase activity whereas the absence of a wide substrate binding pocket in DacC justifies its reduced DD-CPase activity and enhanced β-lactamase activity as β-lactams are smaller molecules than pentapeptides. Moreover, the glutamic acid residue (E167) located at the omega-loop is a strictly conserved amino acid among all the classes of β-lactamases, and plays a key role in the deacylation reaction promoting β-lactam hydrolysis. It activates and brings the hydrolytic water molecule required for deacylation of the acyl enzyme intermediate (Massova et al., 1998; Chowdhury et al., 2012b; Bansal et al., 2015). Hence, the role of glutamic acid is vital for β-lactamase activity. The absence of glutamic acid (E167) in DacD possibly restricts the activation of water molecules to switch on the catalytic interactions, thus preventing the β-lactam hydrolytic activity of DacD. Additionally, the glutamic acid (E75) projecting towards the active site pocket of DacC is believed to play a major role in the β-lactamase activity of the enzyme (Bansal et al., 2015). This residue is substituted by alanine in DacD which further hinders β-lactam hydrolytic activity.

The docking results indicate that the interaction of DacC with the β-lactam substrates is close as compared to DacD explaining its enhanced ability to hydrolyse β-lactam antibiotics. Evaluation of these interactions (bond distances and affinities) along with the docking results (**Fig. S5 a, b and c**) further elucidates the possible mechanisms for the increased β-lactamase activity for DacC. It is well established that the hydrolysis of the β-lactam substrate is accomplished by a nucleophilic attack of the active site deprotonated serine of the SXXK tetrad. We observed that in all the cases of the docking complex of DacC with the β-lactam antibiotic, the active site serine (S58 in DacC and S60 in DacD) is situated in the range of hydrogen bond formation, indicating the probability of a strong interaction, whereas these bond lengths are significantly greater in the docked complexes of DacD, thereby inhibiting proper interactions which is reflected in the distances tabulated in **Table S4.** In the DacC-β-lactam docked complexes the substrates are situated at a closer proximity to the protein (**Table S4**) and mostly interact via H-bond and electrostatic interactions but the same distances drastically increase in the docked complexes of DacD. The increased distances in case of DacD might prohibit the chances of interactions, thus rendering the protein incapable of hydrolysing β-lactam substrates.

The affinity of different PBPs towards different β-lactams is strain specific. Here we witness the differential binding affinity of *A. baumannii* DD-CPases towards different groups of β-lactams. Recently, the IC_50_ values of purified *A. baumannii* PBP1a and PBP3 for different β-lactams and inhibitors were determined. The results indicated that both the PBPs had high affinities for penicillins and cephalosporins. However, DacD was inhibited more efficiently by carbapenems as compared to DacC. Also, β-lactamase inhibitors (clavulanic acid, sulbactam) demonstrated selective inhibition of DacD. Previously it was found that the relative affinity of β-lactamase inhibitors was less for PBP1a as compared to PBP3 (Papp-Wallace et al., 2012).

Overall, it is very interesting to note how DacC, in spite of being an LMM PBP, which is otherwise reported as a dispensable protein in other bacterial species, plays a multifaceted role in this dreadful nosocomial pathogen. A comprehensive understanding of the binding affinities of different antibiotics to their target PBPs might facilitate the development of better combination therapies against *A. baumannii*.

## Conclusion

Based on the obtained results, we conclude that DacD is a mono-functional DD-CPase whereas DacC is a dual enzyme possessing both DD-CPase as well as β-lactamase activities.

## Supporting information

Figure S1, S2, S3, S4, S5; Table S1, S2, S3, S4, S5; Supplementary File Methodology

## Acknowledgements

SP and DJ indebted to IIT Kharagpur for their doctoral fellowship and SB is supported by DST-INSPIRE program her thesis. Authors are grateful to Dr. Bryan W. Davies for gifting us the plasmids pAT03, pAT04, pABBRMCS and pKD4, and for his valuable suggestions regarding knockout construction in *A. baumannii*. We thank Dr. Akash Kumar, Prof. Sudip Kumar Ghosh, Dr. Dhriti Mallik and the members of Molecular Microbiology laboratory, Department of Bioscience and Biotechnology, IIT Kharagpur for their support and cooperation.

## Author contributions

Conception or design of the study: SP, ASG; Acquisition, analysis, or interpretation of the data: SP, DJ, SB, GK, SKR; Writing and editing of the manuscript: SP, DJ, ASG.

