## Supplementary material for "*Acinetobacter baumannii* DacC influences cell shape, biofilm formation and physiological fitness by manifesting DD-carboxypeptidase, and β-lactamase dual-enzyme activities": Figure S1, S2, S3, S4, S5; Table S1, S2, S3, S4, S5; Supplementary File Methodology

The dual activity penicillin-binding protein DacC possessing both DD-carboxypeptidase and β-lactamase activities affects the cell shape, biofilm formation and fitness of *Acinetobacter baumannii*

**Supplementary Figure Legends**

**Figure S1. Genetic Manipulations.**

a. **Confirmation of deletion of *dacC* and *dacD* from AB19606 and AB17904.** Agarose gel electrophoresis of the PCR products using the genomic DNA of the wild type and knockout strains as templates: (i) Single and Double knockouts in AB19606 using *dacC* gene cloning primers: Lane 1: AB19606; Lane 2: AB19606∆C; Lane 3: AB19606∆D; Lane 4: AB19606∆C∆D*.* (ii) Single and Double knockouts in AB19606 using *dacD* gene cloning primers: Lane 1: AB19606; Lane 2: AB19606∆D; Lane 3: AB19606∆C; Lane 4: AB19606∆C∆D*.* M: 1 kb DNA Ladder.
b. **Cloning of *dacC* and *dacD* genes in pBAD18-Cam plasmid. Agarose gel electrophoresis:** Lane 1: Restriction endonuclease (RE) digest of pBAD-C with *Kpn*I; Lane 2: RE digest of pBAD-D with *Nhe*I and *Kpn*I; Lane 3: RE digest of pBAD18-Cam with *Kpn*I; Lane 4: RE digest of pBAD-D with *Nhe*I and *Kpn*I; Lane 5: RE digest of pBAD-D with *Kpn*I; M: 10 kb DNA Ladder.
c. **Cloning of *dacC* and *dacD* genes in pABT. Agarose gel electrophoresis:** Lane 1: RE digest of pABT-C with *Kpn*I; Lane 2: RE digest of pABT-C with *Sac*I and *Kpn*I; Lane 3: RE digest of pABT-D with *Sac*I and *Kpn*I; Lane 1: RE digest of pABT-D with *Kpn*I; M: 10 kb DNA Ladder.
d. **Cloning of *dacC* and *dacD* genes in pET-28(a)+. Agarose gel electrophoresis:** Lane 1: RE digest of pET28 plasmid with *Nhe*I; Lane 2: RE digest of pET-sC with *Nhe*I and *Sac*I; Lane 3: RE digest of pET-sD with *Nhe*I and *Sac*I; M: 10 kb DNA Ladder.

**Figure S2.** Growth rate of A. baumannii reduces in the absence of dacC. Growth analysis of A. baumannii AB19606 (WT) and the DD-CPase knockout mutants (the single knockouts AB19606ΔdacC (ΔC), AB19606ΔdacD (ΔD), and the double knockout AB19606ΔdacCΔdacD (ΔCΔD). Bacteria were grown at 37 °C in LB media in static condition and the OD600 was measured for 21 hrs every 30 mins. The graph is plotted on a semilogarithmic base 2 scale (y axis). The graph is a representative of three independent experiments.

**Figure S3. (a) Purification of soluble proteins.** 15% SDS-PAGE: (i) Expression of sDacC: Lane 1: Uninduced pET-C; Lane 2: Induced pET-C (0.05 mM IPTG); Lane 3: dialysed pure sDacC; (ii) Expression of sDacD: Lane 1: Uninduced pET-D; Lane 2: Induced pET-D (0.5 mM IPTG); Lane 3: dialysed pure sDacD; M: Protein Marker. **(b) MALDI-ToF analysis of the purified PBPs**. (i) MALDI-ToF of sDacC; (ii) MALDI-ToF of sDacD.

**Figure S4. Representative graphs depicting the inhibition of sDacC and sDacD with**

a. **Carbapenems** (i) meropenem MER, (ii) doripenem DOR;
b. **Cephalosporins** (i) cefuroxime CXM, (ii) cefexime CFX, (iii) cefsulodin CFS.

**Figure S5. β-lactam substrate binding to the active site residues of the DacC and DacD proteins**.

a. Docking with piperacillin (representative of penicillin group);
b. Docking with meropenem (representative of carbapenem group);
c. Docking with cefsulodin (representative of cephalosporin group).

[Active site residues are shown in line model and the substrates are represented in stick form. The interacting distances are represented as dotted lines].

**Supplementary Figures**

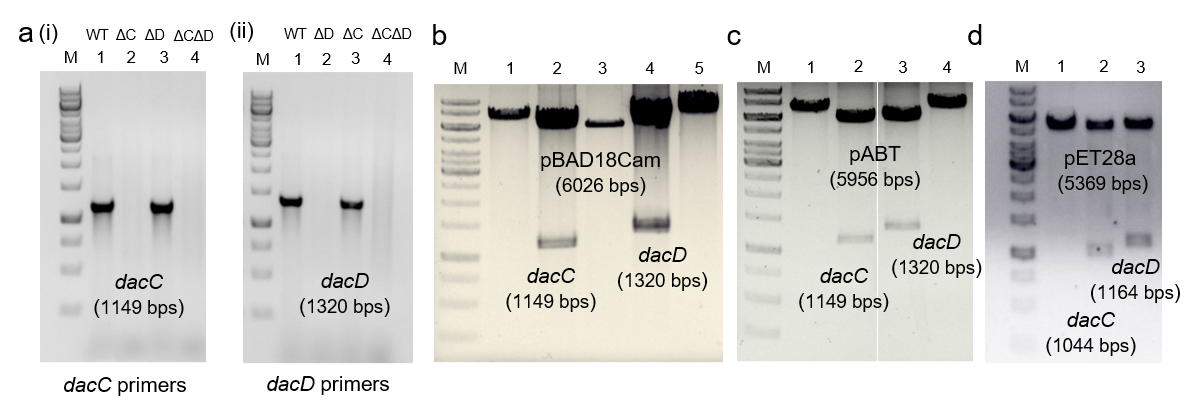

**Figure S1**

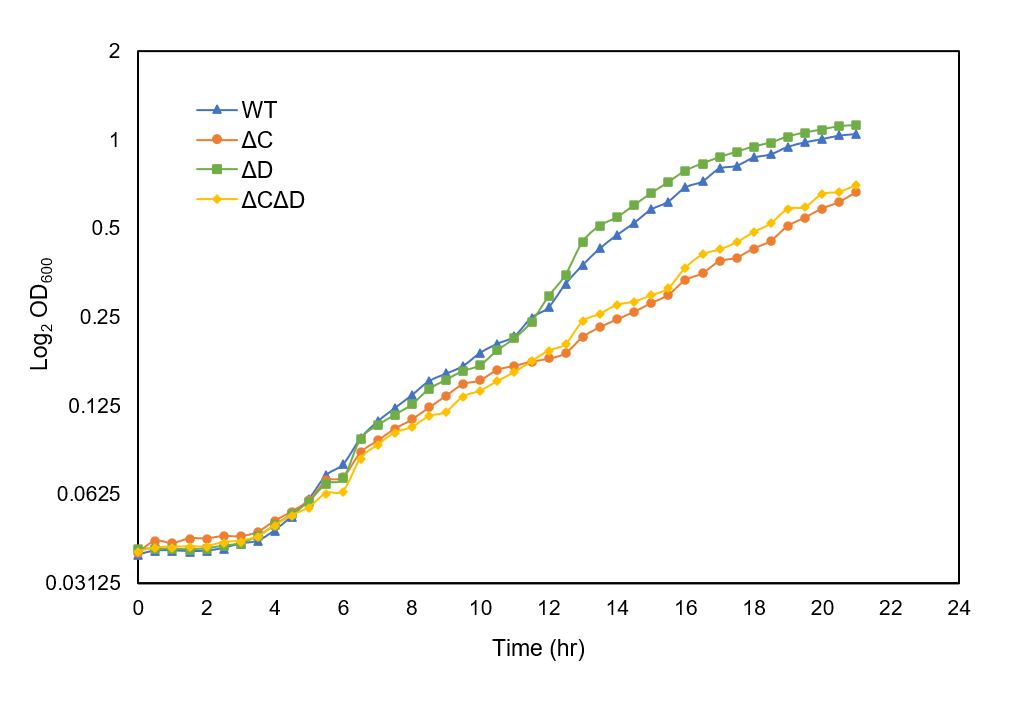

**Figure S2**

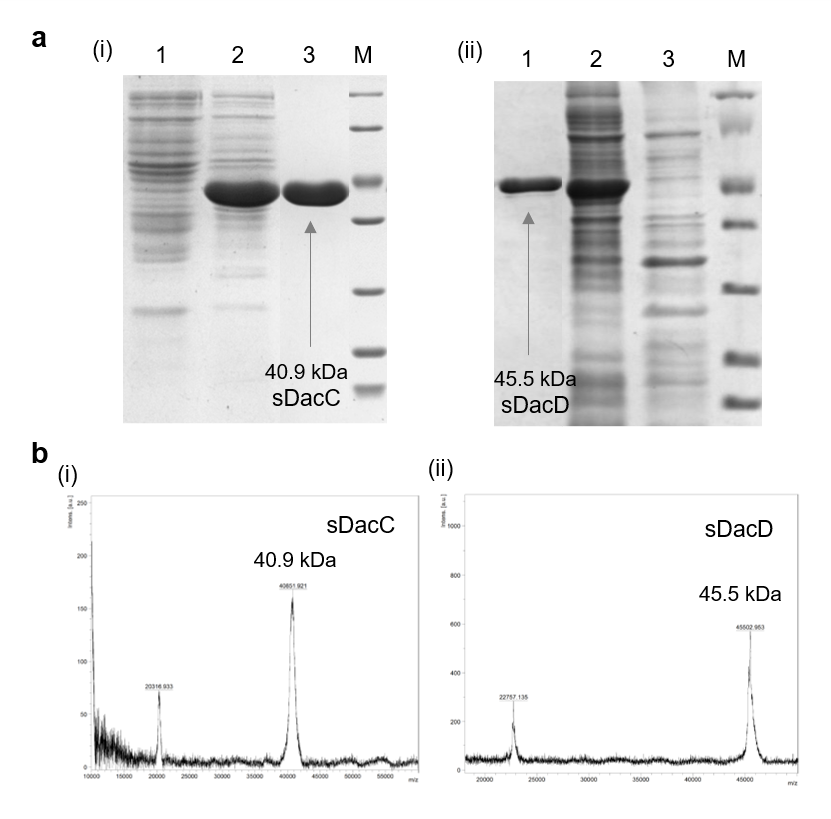

**Figure S3**

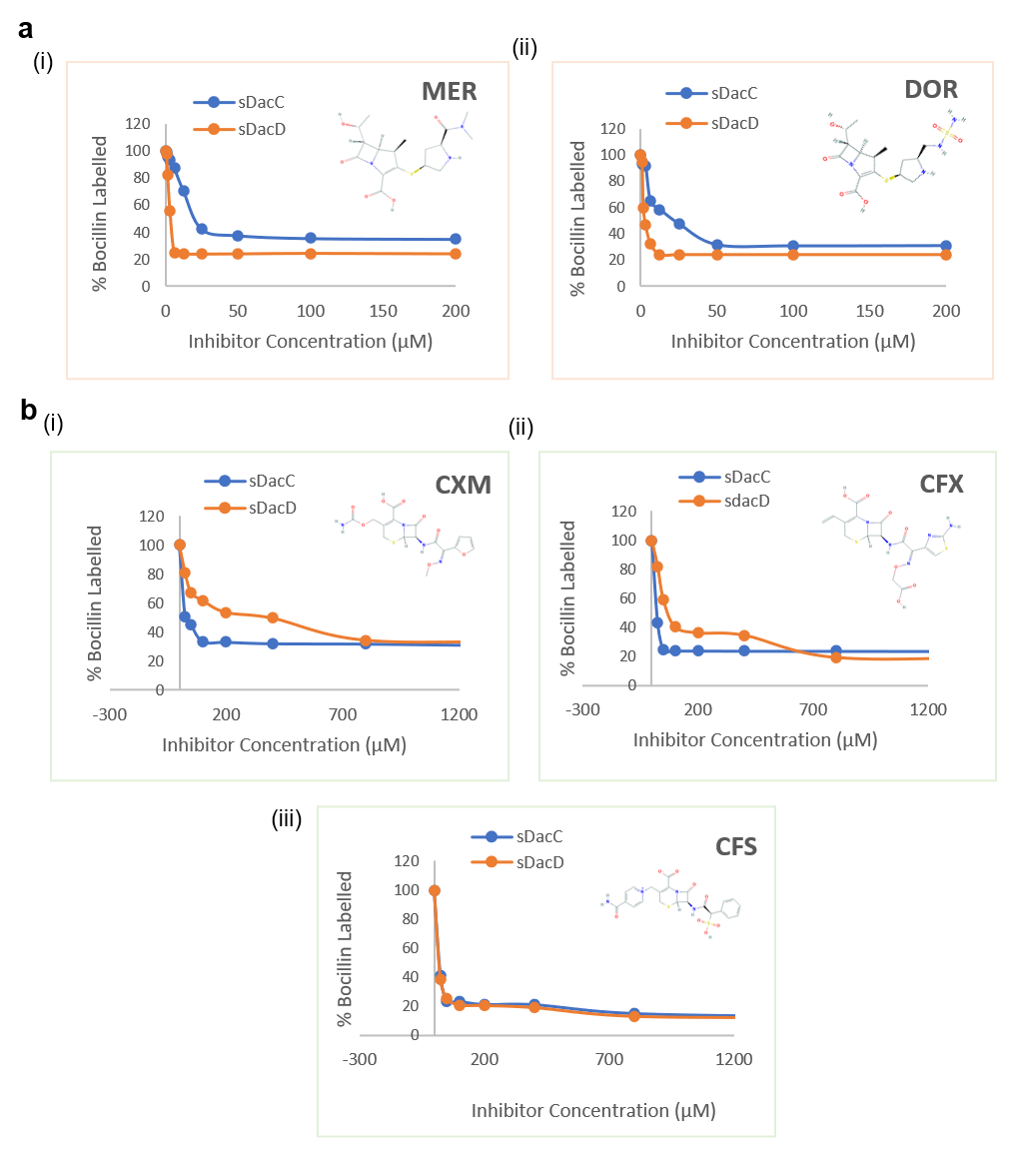
**Figure S4**

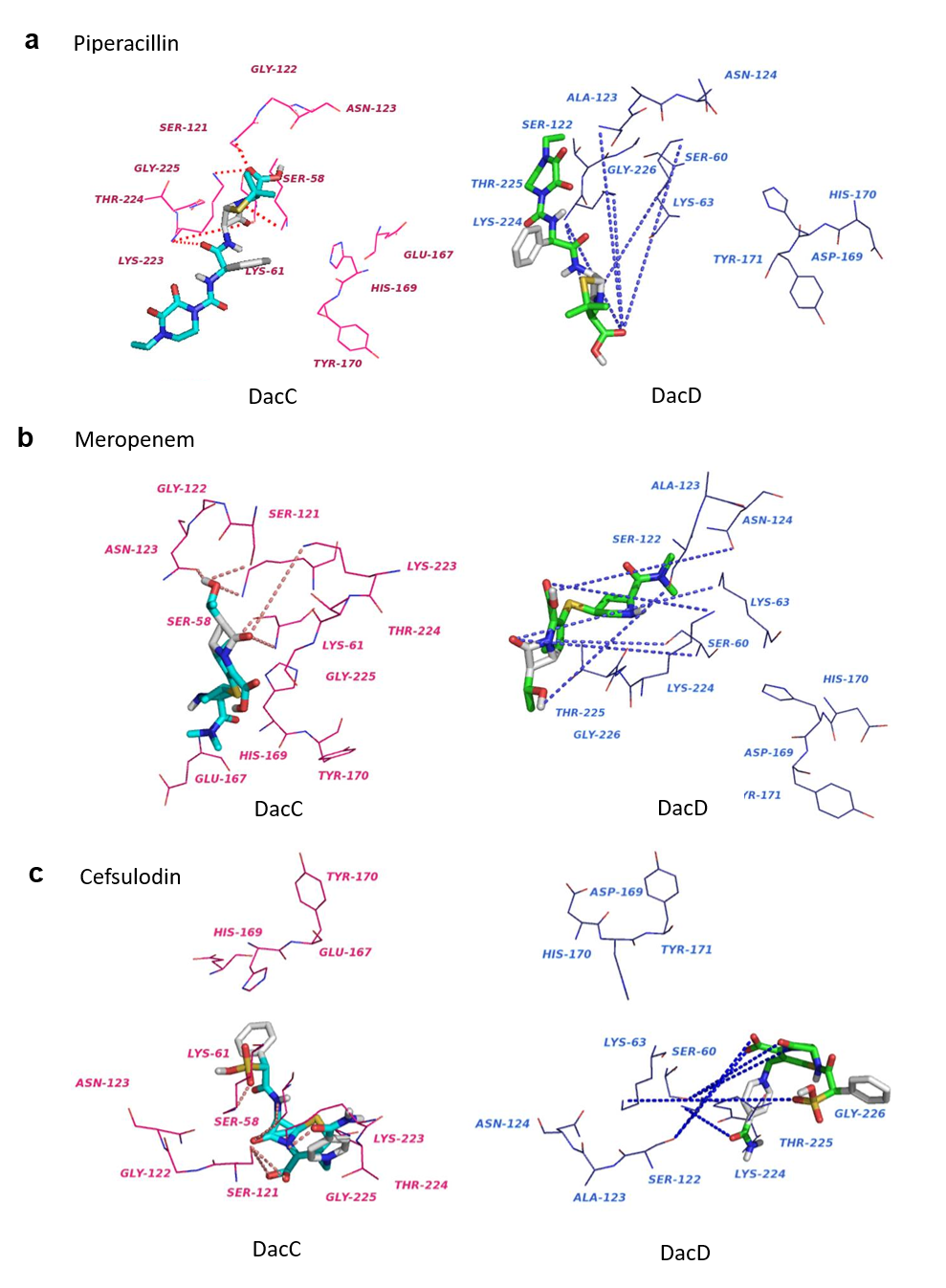

**Figure S5**

**Supplementary Table Legends**

**Table S1. Primer Sequences**

| **Gene** | **Primer Sequence (5’-3’)** |
| --- | --- |
| **Deletion Primers** | |
| dacC_Del_FP | GCTTTATCCTAAAATGCTAGGCTCTTCTGCTTCATACAAGATATTGGAATTACCTAGATGTGTAGGCTGGAGCTGCTTC |
| dacC_Del_RP | CACTCACGCTAATAGCATGAGTGCTTAATTCAACTATTCTAAAATTATTTAAATT TAGAACATATGAATATCCTCCTTAG |
| dacD_Del_FP | CTTCTACAAAAAAAAGGCATAGCATGCTGGGCATCTGTGTTCCGTATTTGCCAGTAAACGTCAAGGTCACCTATGTGTAGGCTGGAGCTGCTTC |
| dacD_Del_RP | GATGGCTGAATTATGCCATCTTTTCTTTATAAAATTTATTGTGCATCTGTGCACAATTTGATGTTTCATATGAATATCCTCCTTAG |
| **Cloning Primers (pBAD18-Cam)** | |
| dacC_BAD_FP | CTCTATGCTAGCAGGAGGATCTATCTATGACTCGAAAAAGCGC |
| dacC_BAD_RP | CTCTCTGGTACCTTAGAATAAGTTGCTGAAGAATTGTTTGATATGG |
| dacD_BAD_FP | CTCTCTGAGCTCAGGAGGCTCTCTCTGTGAAATTCTTCCTATCTC |
| dacD_BAD_RP | CTCTCTGGTACCTTAGTGCGAATCTATAGGG |
| **Cloning Primers (pABT)** | |
| dacC_ABT_FP | CTCTATGGTACCAGGAGGATCTATCTATGACTCGAAAAAGCGC |
| dacC_ABT_RP | CTCTCTGAGCTCTTAGAATAAGTTGCTGAAGAATTGTTTGATATGG |
| dacD_ABT_FP | CTCTCTGGTACCAGGAGGCTCTCTCTGTGAAATTCTTCCTATCTC |
| dacD_ABT_RP | CTCTCTGAGCTCTTAGTGCGAATCTATAGGG |
| **Cloning Primers (pET28a)** | |
| sdacC_PET_FP | CTCTCTGCTAGCGCAACTGTCTTATCAGCTCCG |
| sdacC_PET_RP | CTCTCTGAGCTCTTATTAGAAGAAACCAGCTTCTTCAACAGG |
| sdacD_PET_FP | CTCTCTGCTAGCGCTCTACTTAATATTGCTCCTG |
| sdacD_PET_RP | CGCTCGGAGCTCTCAGAAAATATTTGCTTCTTCAATGTG |
| **qPCR Primers** |  |
| dacC_qPCR_FP | GAACGTGCTGACCAAACTCG |
| dacC_qPCR_RP | GAGTTTTAATGCCGTCCGC |
| dacD_qPCR_FP | TAGGTACACCAAGTGCCGTG |
| dacD_qPCR_RP | TCGTAACAATGGTTGGCTGC |
| 16SrRNA_qPCR_FP | TGCTCAAGTCCCAGCGTAAC |
| 16SrRNA_qPCR_RP | CCAGCAAAACTTCCGGTTC |

**Table S2. Experimental conditions for determination of kinetic parameters**

| **β-Lactam Group** | **Antibiotic** | **Concentration range**  (µM) | **Molar extinction coefficient**  Δɛ (M^-1^cm^-1^) | **Wavelength**  λ (nm) | **Enzyme concentration used** (µg) |
| --- | --- | --- | --- | --- | --- |
| **Penicillin** | **Penicillin G** | 500-2000 | -936 | 235 | 0.5-1 |
|  | **Ticarcillin** | 500-2000 | -673 | 235 | 0.5-1 |
|  | **Piperacillin** | 500-2000 | -936 | 232 | 0.5-1 |
| **Cephalosporin** | **Cefotaxime** | 50-100 | -7250 | 264 | 0.5-1 |
|  | **Cefsulodin** | 50-100 | -8990 | 260 | 0.5-1 |
|  | **Ceftazidime** | 50-120 | -10300 | 265 | 0.5-1 |
| **Carbapenem** | **Doripenem** | 10-50 | -11500 | 295 | 0.5-1 |
|  | **Meropenem** | 10-50 | -6500 | 300 | 0.5-1 |
|  | **Imipenem** | 10-50 | -9000 | 300 | 0.5-1 |

**Table S3. β-lactam susceptibility of *E. coli* Δ*dacA* mutants expressing *A. baumannii* DD-CPases**

| **Strains** | **PIP** | **AMX** | **CFR** | **CEF** |
| --- | --- | --- | --- | --- |
| CS109 | 2 | 8 | 32 | 16 |
| AM15- pBAD18-Cam | 0.5 | 2 | 8 | 2 |
| AM15-pPJ5 | 2 | 8 | 32 | 16 |
| AM15- pBAD-C | 2 | 8 | 32 | 16 |
| AM15-pBAD-D | 0.5 | 2 | 4 | 2 |

[PIP: Piperacillin; AMX: Amoxicillin; CFR: Cefadroxil; CEF: Cephalothin] **Table S4. Percentage distribution of secondary structures of the purified PBPs**

|  | **CD spectroscopy** | | | | **PREDICT PROTEIN** | | |
| --- | --- | --- | --- | --- | --- | --- | --- |
|  | α-Helix | β-Sheet | Turn | Random coil | Helix | Strand | Loop |
| **sDacC** | 27.5 | 13.3 | 32.4 | 26.8 | 30.1 | 21.47 | 48.43 |
| **sDacD** | 24.3 | 8.7 | 30.6 | 36.4 | 29.38 | 18.68 | 51.94 |

**Table S5. Interacting bond lengths of the PBP- substrate docked complexes**

|  | **Interacting atoms** | **DacC**  **(Interacting distance)** | **DacD**  **(Interacting distance)** |
| --- | --- | --- | --- |
| **Piperacillin** | Ser 58/60 -β-lactam ring ‘N’ | 2.69 Å | 9.85 Å |
|  | Lys61/63 - (β-lactam ring ‘N’) & (β-lactam ring ‘O’) | 3.07 Å & 2.99 Å | 14.19 Å & 7.46 Å |
|  | Ser 121 – ‘O 31’ | 2.43 Å | 13.20 Å |
|  | Lys 223 – β-lactam ring ‘O’ & O ‘32’ | 2.99 Å & 3.93 Å | 9.75 Å & 11.12 Å |
| **Meropenem** | Ser 58/60 -β-lactam ring ‘N’ & β-lactam ring ‘O’ | 3.05 Å & 3.17 Å | 7.26 Å & 12.07 Å |
|  | Lys 61/63 - β-lactam ring ‘O’ | 2.65 Å | 13.41 Å |
|  | Ser 121 – ‘C28’ | 3.28 Å | 10.51 Å |
|  | Lys 223 – ‘β-lactam ring ‘O’ | 4.47 Å | 11.16 Å |
|  | Asn 123 – ‘C28’ | 1.81 Å | 12.76 Å |
| **Cefsulodin** | Ser 58/60 -β-lactam ring ‘N’ | 2.95 Å | 9.81 Å |
|  | Lys 61/63 - β-lactam ring ‘N’ & β-lactam ring ‘O’ | 3.24 Å & 2.69 Å | 9.19 Å & 13.62 Å |
|  | Ser 121 – ‘O3 & O4’ | 3.87 Å & 4.01 Å | 9.33 Å & 8.48 Å |
|  | Lys 223/224 – β-lactam ring ‘O’ | 2.35 Å | 6.23 Å |

**Supplementary File Methodology**

Molecular modelling & bioinformatic analysis

Homology modelling relies on the identification of two or more template structures which are likely to resemble the target or query sequence. The 3D structural models of both DacC and DacD were prepared using a restraint-based modelling program known as MODELLER v9.16 (Sali & Blundell, 1993). The optimization of the target-template alignment, comparison between root mean square (RMS), distance root mean square (DRMS), atomic distances, and dihedral angle corrections were done at the backend of the programme to generate several tertiary models. The best structural models for both the proteins were chosen on the basis of the lowest discrete optimized protein energy (DOPE) score and retained for further structural refinement. Subsequently, the predicted 3D models were also validated using PROCHECK and found to be stereo-chemically stable in the Ramachandran plot. Afterwards the models were subjected to an energy refinement process for quality improvement through the YASARA minimization server using atomic AMBER force field (Krieger *et al.*, 2009). Finally, the refined stereo-chemically stable and energy optimised 3D models were obtained and taken for further investigation for legitimacy.

For docking analysis, the Autodock4.2 software was used (Morris *et al.*, 2009). The grid box was generated around the active site by considering appropriate x, y and z coordinate positions of the protein and Kollman charges were assigned before generating the grid parameter file. The docking parameters were set using the Lamarckian genetic algorithm to generate a dock parameter file and the final docking was performed by the AutoDock module present in the AutoDock4.2 suite. Finally, the lowest energy docked conformations of the ligands were selected based on the scoring function (Z-score) of docking with the optimized models of the proteins and the results were analysed to assess the variation in DD-CPase and beta-lactamase activities.

**Circular Dichroism**

The protein secondary structure analysis was done using circular dichroism (CD) spectral analysis with JASCO J-815 CD spectrometer (JASCO international company limited, Tokyo, Japan) at 190–240 nm wavelengths, temperature 37°C; data were recorded in a 0.1 cm pathlength cuvette with 1 mg mL^−1^ protein concentration in 10 mM Tris-Cl buffer having 150 mM NaCl (pH 7.4). Data were recorded at 0.2 nm step resolution; time constant 1 s, sensitivity 10 milli degrees, with scan speed of 50 nm min^−1^ and a spectral bandwidth of 2 nm. To reduce the noise and random instrumental error, an average spectrum of four successive accumulations were taken over 190–240 nm range.

**MALDI- ToF**

The molecular mass of the purified proteins was determined by MALDI-ToF spectral analysis was done using a VOYAGER- DE PRO, Applied Biosystem, USA.
